# Specific targeting of inflammatory osteoclastogenesis by the probiotic yeast *S. boulardii* CNCM I-745 reduces bone loss in osteoporosis

**DOI:** 10.1101/2022.08.01.502322

**Authors:** Maria-Bernadette Madel, Julia Halper, Lidia Ibáñez, Claire Lozano, Matthieu Rouleau, Antoine Boutin, Rodolphe Pontier-Bres, Thomas Ciucci, Majlinda Topi, Christophe Hue, Jérôme Amiaud, Salvador Iborra, David Sancho, Dominique Heymann, Henri-Jean Garchon, Dorota Czerucka, Florence Apparailly, Isabelle Duroux-Richard, Abdelilah Wakkach, Claudine Blin-Wakkach

## Abstract

Bone destruction is a hallmark of chronic inflammation, and bone-resorbing osteoclasts arising under such a condition differ from steady-state ones. However, osteoclast diversity remains poorly explored. Here, we combined transcriptomic profiling, differentiation assays and *in vivo* analysis in mouse to decipher specific traits for inflammatory and steady state osteoclasts. We identified and validated the pattern-recognition receptors (PRR) TLR2, Dectin-1 and Mincle, all involved in yeast recognition as major regulators of inflammatory osteoclasts. We showed that administration of the yeast probiotic *Saccharomyces boulardi* CNCM I-745 (*Sb*) *in vivo* reduced bone loss in OVX but not sham mice by reducing inflammatory osteoclasts. This beneficial impact of *Sb* is mediated by the regulation of the inflammatory environment required for the generation of inflammatory osteoclasts. We also showed that *Sb* derivatives as well as agonists of TLR2, Dectin-1 and Mincle specifically inhibited directly the differentiation of inflammatory but not steady state osteoclasts *in vitro*. These findings demonstrate a preferential use of the PRR-associated costimulatory differentiation pathway by inflammatory osteoclasts, thus enabling their specific inhibition, which opens new therapeutic perspectives for inflammatory bone loss.

## Introduction

Osteoclasts (OCLs) are multinucleated phagocytes derived from monocytic progenitors and specialized in bone resorption (Madel et al., 2019). Similar to other cells of monocytic origin, they are also innate immunocompetent cells and eterogeneous in their phenotype, function and origin (Ibáñez et al., 2016; Jacome-Galarza et al., 2019; Madel et al., 2020, 2019; Yahara et al., 2020). OCLs derived from steady state bone marrow (BM) cells or from BM CD11b^+^ monocytic cells (MN-OCLs) promote tolerance by inducing CD4^+^ and CD8^+^ regulatory T cells (tolerogenic OCLs/t-OCLs) (Ibáñez et al., 2016; Kiesel et al., 2009). In contrast, in the context of bone destruction linked to inflammatory bowel disease (IBD) or when derived from dendritic cells (DC-OCLs), OCLs induce TNFα-producing CD4^+^ T cells (inflammatory OCLs/i-OCLs) (Ibáñez et al., 2016; Madel et al., 2020). OCLs associated with inflammation can be identified by expression of Cx3cr1 (fractalkine receptor) and the proportion of Cx3cr1^+^ OCLs increases in osteoporosis, IBD and after RANK-L treatment (Ibáñez et al., 2016; Madel et al., 2020). However, Cx3cr1 is only expressed in approximately 20% of i-OCLs (Ibáñez et al., 2016; Madel et al., 2020), which highlights their heterogeneity while limiting the possibility to analyze them in the context of pathological bone loss and urges the identification of novel markers.

Current anti-resorptive therapies aim to globally inhibit OCLs, without considering their recently established diversity. In the long term, they result in poor bone remodeling which may increase the risk of atypical fractures (Reyes et al., 2016). Therefore, an in-depth characterization of OCLs associated with healthy versus inflammatory bone resorption would allow the identification of distinct characteristics that could help to specifically target i-OCL.

Inflammatory OCLs arise under the control of persistent high levels of RANK-L, IL-17 and TNFα mainly produced by CD4^+^ T cells that play a major role in pathological osteoclastogenesis observed in osteoporosis and IBD (Cenci et al., 2000; Ciucci et al., 2015; Ibáñez et al., 2016; Li et al., 2011). Interestingly, the emergence of such osteoclastogenic CD4^+^ T cells is associated with gut dysbiosis and increased intestinal permeability (Jones et al., 2017; Li et al., 2016). In line with this, bacterial probiotics such as *Lactobacillus* and *Bifidobacteria* have been shown to effectively reduce osteoporotic bone loss (Li et al., 2016; Britton et al., 2014; Ohlsson et al., 2014; Ibáñez et al., 2019b), but their specific effect on inflammatory OCLs remains unknown.

Here, using a comparative RNAseq approach performed on sorted pure mature MN-OCLs and DC-OCLs as models of t-OCLs and i-OCLs, respectively, as already established (Ibáñez et al., 2016; Madel et al., 2020), we showed that the two OCL populations are distinctly equipped to respond to different signals that can modulate their differentiation. In particular, the pattern recognition receptors (PRRs) Dectin-1, TLR2 and Mincle involved in the response to fungi (Li et al., 2019; Sancho and Reis e Sousa, 2012) are overexpressed in i-OCLs.

Administration of the probiotic yeast *Saccharomyces boulardii* CNCM I-745 (*Sb*) which is used in the treatment of gastrointestinal disorders for its anti-inflammatory properties and its capacity to restore the gut microbiota (Czerucka and Rampal, 2019; Terciolo et al., 2019) significantly reduces bone loss and inflammatory parameters *in vivo* in ovariectomized (OVX) mice. *In vitro, Sb* derivates as well as agonists of the PRRs overexpressed in i-OCLs specifically inhibit the differentiation of these cells without affecting t-OCLs. These data open perspectives on targeting specific OCL populations and provide evidence for the protective effect of a probiotic yeast on inflammatory bone resorption. Our study unveils very new insights into the regulation and modulation of i-OCLs and enables a better understanding of the molecular mechanisms involved in inflammation-induced bone erosions.

## Results

### Transcriptomic profiling reveals upregulation of innate immune receptors in i-OCLs

To better understand the differences between t-OCLs and i-OCLs, we performed a comparative RNA-sequencing (RNAseq) approach between sorted mature (≥3 nuclei) MN-OCLs (originating from BM CD11b^+^ monocytic cells) and DC-OCLs (differentiated from BM-derived DCs) and representing t-OCLs and i-OCLs, respectively, as already demonstrated (Ibáñez et al., 2016; Madel et al., 2020) (Supp: Figure 1a-b). A total of 906 genes (Log_2_FC≥1; p<0.05) were significantly differentially expressed between both OCL subsets (Figure. 1a), including Cx3cr1 previously identified as a marker of i-OCLs (Ibáñez et al., 2016; Madel et al., 2020). The most differentially expressed genes were related to innate immunity and immune defense responses (Figure 1b-c), confirming our previous observation that i-OCLs and t-OCLs differ in their immune capacity (Ibáñez et al., 2016).

**Figure 1:**
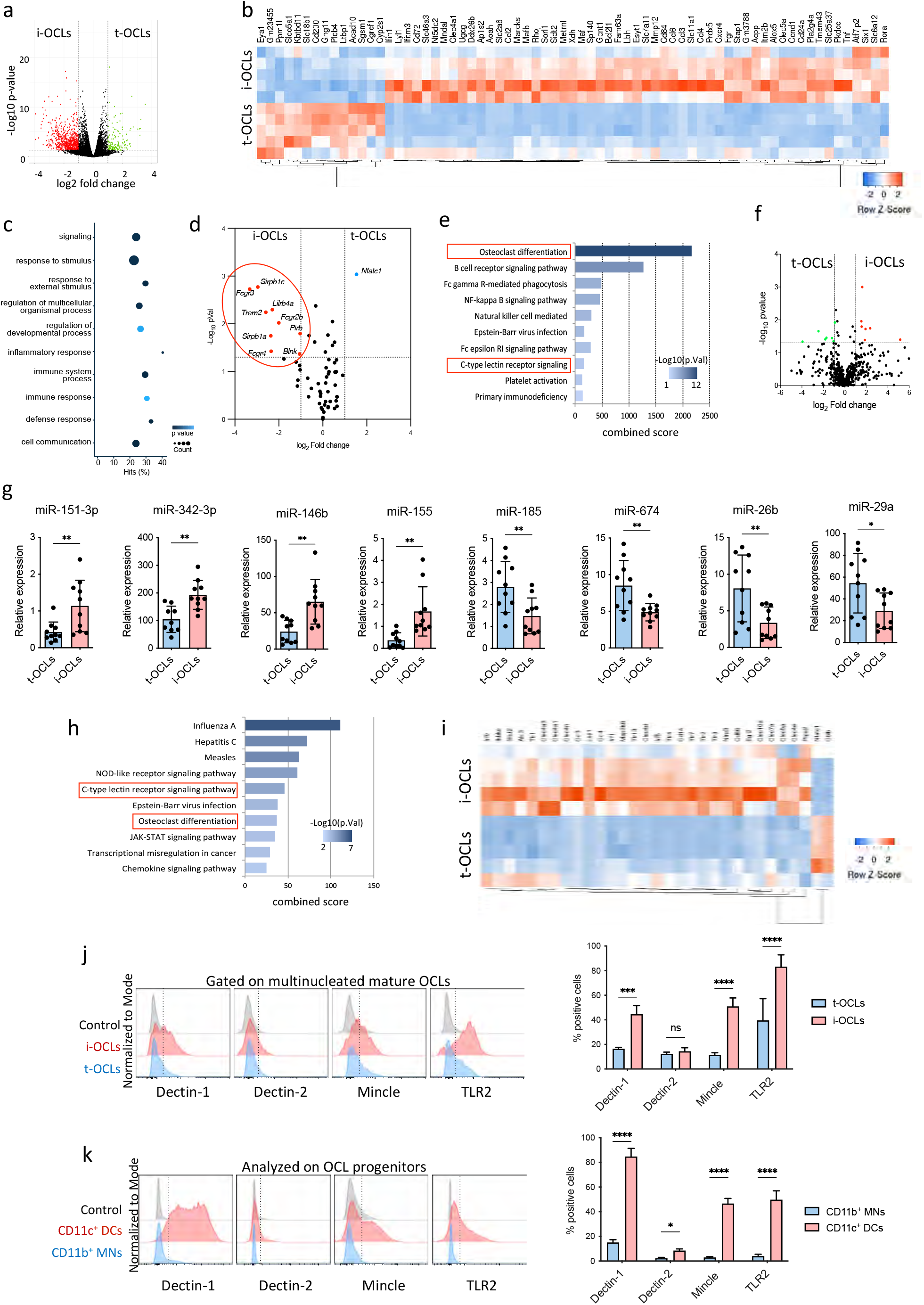
Comparative transcriptomic analysis of i-OCLs vs. t-OCLs reveals two distinct populations of OCLs and differences in their differentiation pathways. **(a)** Volcano plot analysis of differentially expressed genes between i-OCLs (DC-derived OCLs) and t-OCLs (MN-derived OCLs) (p<0.05; FC≥2) performed on n=5 biological replicates for each group. **(b)** Heatmap visualization of the z-scored expression of the top 70 genes (p<0.05; FC≥2) that are significantly differentially expressed between the two OCL subsets. (**c)** Gene ontology analysis of biological processes associated with differentially expressed genes. (**d)** Transcriptomic analysis of selected genes (from Kegg mmu04380, osteoclast differentiation) involved in bone resorption (*Acp5, Car2, Clcn7, Ctsk, Mmp9, Ostm1, Tcirg1)*, the RANK differentiation pathway (*Chuk, Ikbkb, Ikbkg, Map2k1, Map2k7, Map3k7, Mapk8, Mapk9, Nfatc1, Nfkb1, Nfkb2, Tab1, Tab2, Tnfrsf11a, Traf2, Traf6*) and the co-stimulatory differentiation pathway (*Blnk, Fcgr1, Fcgr2b, Fcgr3, Fcgr4, Lilrb4a, Oscar, Plcg2, Sirpb1b, Sirpb1c, Syk, Trem2, Tyrobp)* in i-OCLs and t-OCLs on n=5 biological replicates per group. **(e)** EnrichR annotation (Kegg) for the genes involved in the OCL co-stimulatory differentiation pathway (from Figure 1d). (**f)** Volcano plot visualization of the comparative miRNome analysis of i-OCLs and t-OCLs performed on n=5 biological replicates for each group. **(g)** RT-qPCR analysis on t-OCLs and i-OCLs (n=10 biological replicates per group). miRNA expression was normalized to the sno202 expression using the 2^-(^Δ^Ct)^ method. (**h)** EnrichR annotation (Kegg) for the target genes of the differentially expressed miRNAs. (**i)** Heatmap visualization of the RNAseq data for selected genes involved in the CLR and TLR signaling pathway (from Kegg) in i-OCLs and t-OCLs. (**j)** FACS plots and quantification of mature i-OCLs and t-OCLs for the expression of Dectin-1 and 2, TLR2 and Mincle. OCLs were gated as shown in Supp. Figure 1b. **(k)** FACS analysis on BM-derived CD11c^+^ DCs, and CD11b^+^ BM cells (used as OCL progenitors) and quantification of positive cells for the expression of Dectin-1 and 2, TLR2 and Mincle each marker. *p<0.05; **p<0.01; ***p<0.001; ****p<0.0001; n.s. non-significant differences.

Volcano plot representation showed that the expression of genes involved in OCL resorbing activity as well as in the RANK-RANKL differentiation pathway was not significantly different in the 2 OCL populations, except for *Nfatc1*. However, major differences were found in genes associated with the Ig-like receptor-dependent costimulatory OCL differentiation pathway (Koga et al., 2004; Merck et al., 2004) (Figure 1d). Interestingly, annotation of these genes (EnrichR/Kegg analysis) highlighted that they are also linked to pathways related to immune responses and to PRRs, notably the C-type lectin receptors (CLRs) (Figure 1e).

These results were confirmed by a global miRNA profiling on sorted mature OCL populations (≥3 nuclei). Volcano plot visualization confirmed that the two OCL subsets differ also in their miRNA expression pattern (Figure 1f). Differentially expressed miRNAs (FC≥2, p<0.05, qPCR cycle threshold (CT) values <32) were further validated by RT-qPCR confirming that miR-151-3p, miR-342-3p, miR-146b and miR-155 were significantly upregulated in i-OCLs, and that miR-185, miR-674, miR-26b and miR-29a were upregulated in t-OCLs (Figure 1g). Using the miRWalk software that provides information on miRNA-target interactions (Dweep and Gretz, 2015), we carried out an integrative analysis of these miRNAs and genes that were significantly differentially expressed between both OCL subsets to find possible relationships. In agreement with the RNAseq analysis, computational analysis of these related genes (EnrichR/ Kegg analysis) also revealed an association with the OCL differentiation pathway as well as the C-lectin-like receptor pathway (Figure 1h).

These data strongly suggest that CLRs could play an important role in the specific properties of i-OCLs and t-OCLs. Analysis of the RNAseq data for the expression of genes involved in the CLRs and TLRs signaling pathways confirmed major differences between t-OCLs and i-OCLs (Figure 1i). Validation by flow cytometry analysis of mature multinucleated OCLs confirmed that the proportion of OCLs expressing Dectin-1, Mincle and TLR2, but not Dectin-2, was significantly higher in i-OCLs compared to t-OCLs (Figure 1j). We also analyzed their expression in OCL progenitors. The proportion of cells expressing Dectin-1, Mincle and TLR2 was much higher in CD11c^+^ BM-derived DCs than in BM CD11b^+^ MNs, and the proportion of i-OCL progenitors expressing Dectin-2 was lower than for the other markers (Figure 1k).

### The probiotic yeast *Saccharomyces boulardii* CNCM I-745 protects from osteoporosis-induced bone loss *in vivo*

The in here investigated OCL populations were derived from purified progenitor cells which is a powerful approach to identify major differences (Ibáñez et al., 2016; Madel et al., 2020) while *in vivo*, OCLs arise from a mixture of different BM progenitors whose proportions are depending on the pathophysiological conditions (Madel et al., 2019). Thus, we compared BM-derived OCLs from OVX-induced osteoporotic mice (OVX) in which the proportion of i-OCL increases (Madel et al., 2020) to SHAM OCLs. FACS analysis revealed higher proportions of Dectin-1^+^ and TLR2^+^ OCLs, in OCLs generated from OVX mice compared to those from SHAM mice, while the proportion of Mincle^+^ OCLs was not significantly altered (Figure 2a).

**Figure 2:**
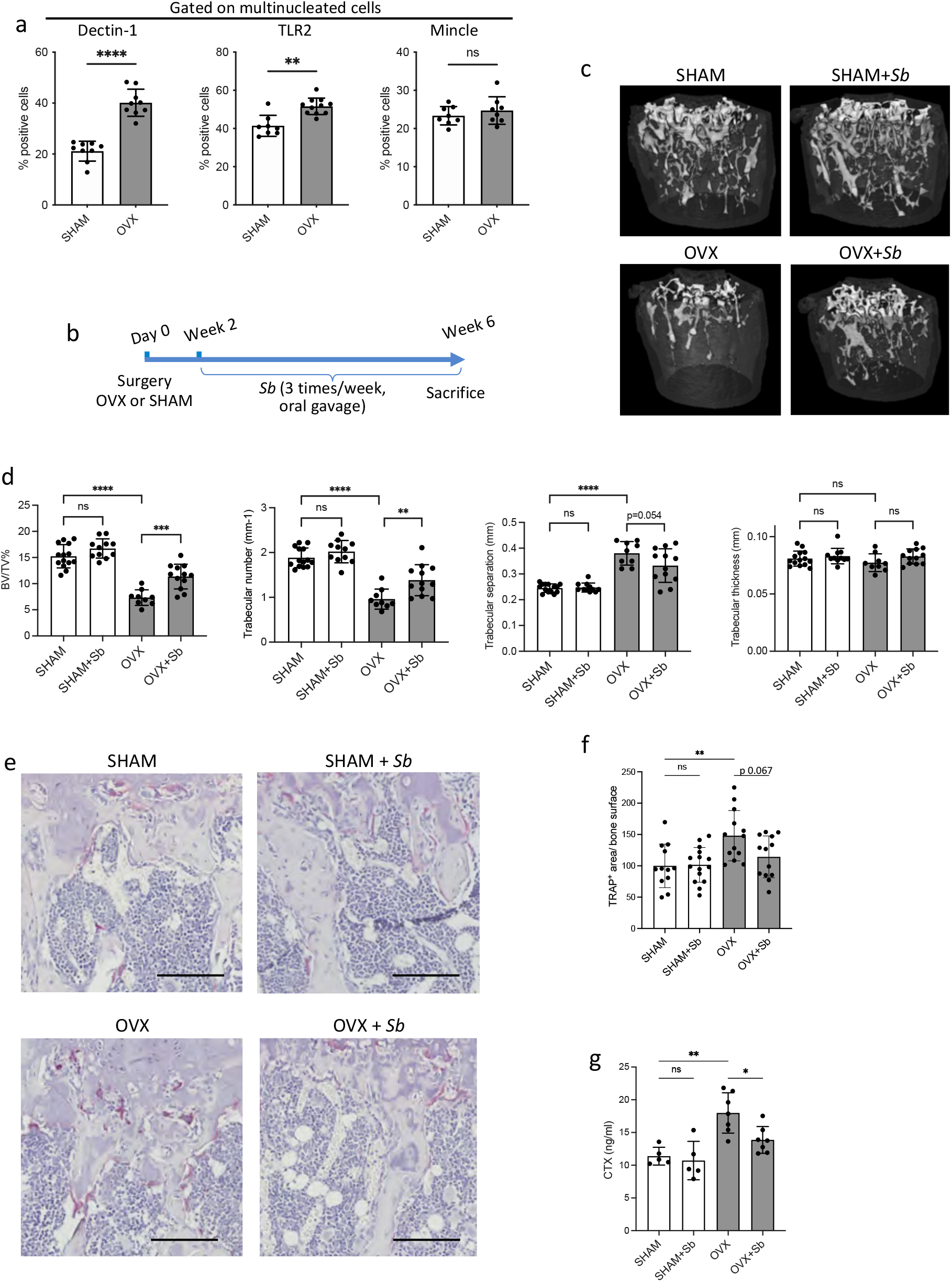
The probiotic yeast *Sb* has beneficial effects on bone loss in osteoporosis. (**a**) Quantification of FACS analysis of Dectin-1^+^, TLR2^+^ and Mincle^+^ mature OCLs (≥3 nuclei, gated as shown in Supp. Figure 1b.) differentiated from the BM of SHAM and OVX mice, 6 weeks after surgery. **(b)** Schematic representation of the experimental procedure. **(c)** Representative μCT images of femurs from SHAM and OVX mice ± *Sb* administration. (**d)** Histograms indicate mean ± S.D. of bone volume fraction (BV/TV), trabecular number, separation and thickness. (**e)** Histological analysis of OCLs using TRAcP staining (in purple) on tibias from SHAM and OVX mice treated or not with *Sb*. Scale bars: 100μm. (**f)** Histogram indicates the mean ± SD of TRAcP^+^ area per bone surface for each condition. Three images of 4-5 biological replicates were analyzed. (**g)** Serum cross-linked C-telopeptides of type I collagen (CTX) were measured by ELISA (n=5 biological replicates per condition). *p<0.05; **p<0.01; ***p<0.001; ****p<0.0001; n.s. non-significant differences.

These PRRs share the capacity to sense fungi and to control responses to pathogenic or commensal gut mycobiome (Li et al., 2019; Sancho and Reis e Sousa, 2012). Therefore, we treated SHAM and OVX mice by oral gavage with the probiotic yeast *Sb* (Figure 2b). As expected, OVX mice showed atrophy of the uterus and their body weight increased compared to SHAM control mice, which was not affected by treatment with *Sb* (Suppl. Figure 2a-b). Micro-computed tomographic (micro-CT) analysis revealed that *Sb*-treated OVX mice displayed reduced bone loss, as assessed by a significantly higher BV/TV and trabecular number and less significant reduction in trabecular separation compared to untreated OVX mice (Figure 2c-d). Furthermore, administration of *Sb* resulted in a reduction of the number of TRAcP^+^ OCLs per bone surface in OVX mice (Figure 2e-f) and a significantly reduced serum level of cross-linked C-telopeptides of type I collagen (CTX), confirming a decrease in OCL activity (Figure 2g). Importantly, *Sb* treatment did not affect cortical parameters (Suppl. Figure 3a). It had no effect on the number of osteoblasts and osteocytes *in vivo* (Suppl. Fig 3b-c) and on the mineralization capacity of osteoblasts *in vitro* (Suppl. Fig 3d).

**Figure 3:**
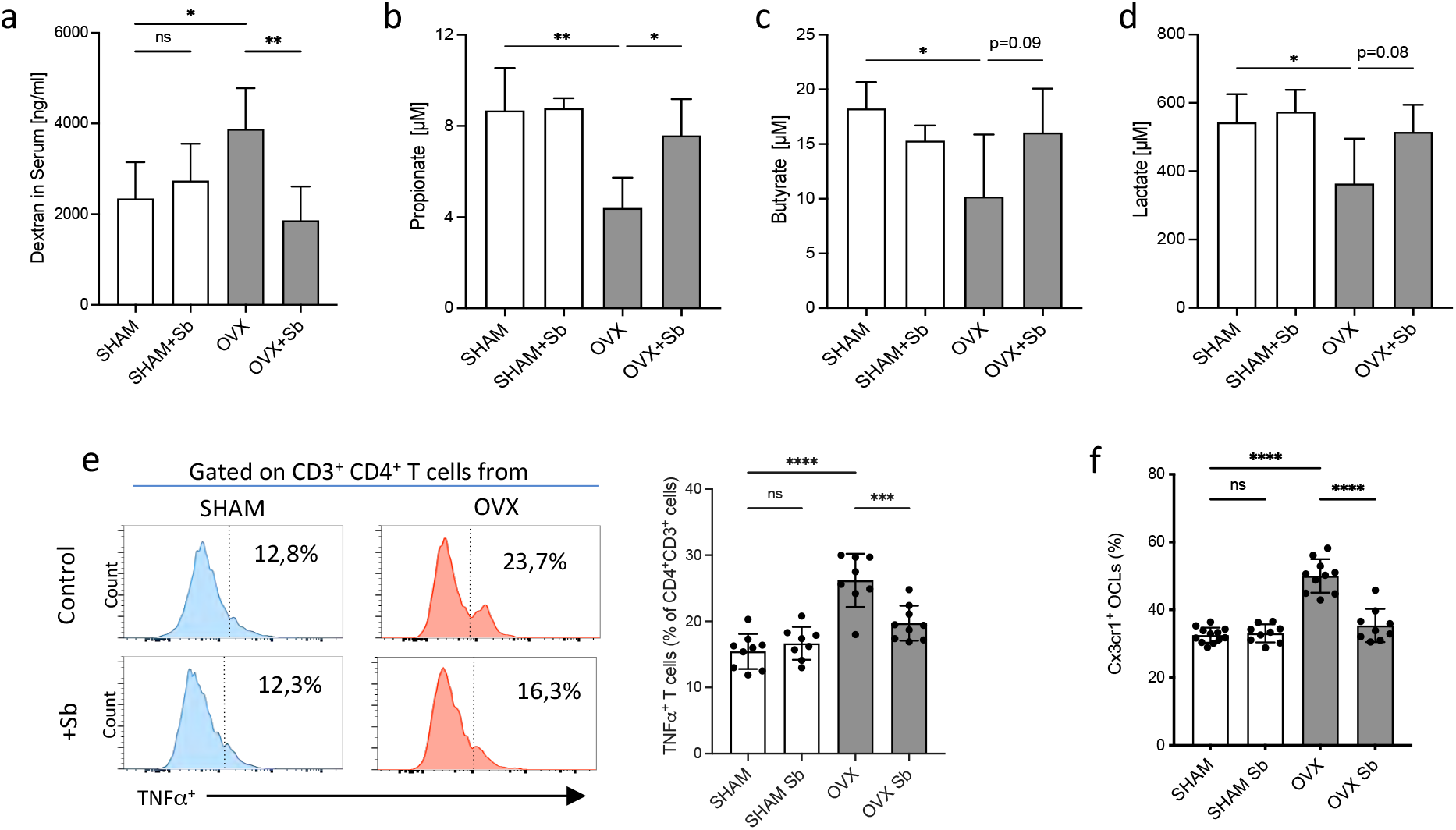
*Sb* reduces inflammatory parameters in osteoporosis. (**a**) Integrity of the intestinal barrier permeability was analyzed by fluorometry on the serum of mice that received oral gavage of dextran-FITC 1 h before sacrifice. (**b-d**) Concentrations of (**b**) propionate, (**c**) butyrate and (**d**) lactate were measured by chromatography in the serum of SHAM and OVX mice treated or not with Sb (n=4 biological replicate per group). *p<0.05; **p<0.01. (**e)** FACS analysis of TNFα-producing CD4^+^ T cells in the BM of OVX and SHAM control mice with or without *Sb* treatment (n=8-10 biological replicate per group). (**f**) Proportion of mature Cx3cr1^+^ BM-derived OCLs (≥3 nuclei, gated as shown in Supp. Figure 1b) from SHAM and OVX mice treated or not with *Sb* (n=10-14 biological replicate per group) was determined by FACS analysis. *p<0.05; **p<0.01; ***p<0.001; ****p<0.0001; n.s. non-significant differences.

We then checked whether *Sb* treatment affects inflammatory parameters and microbiota metabolites that are known to influence bone remodeling (Zaiss et al., 2019). Evaluation of the gut barrier integrity by FITC-dextran assay showed a reduction in serum dextran concentration in *Sb-*treated OVX mice to the level observed in SHAM mice (Figure 3a) confirming the protective effect of *Sb* on the intestinal barrier as already reported in other pathological contexts (Terciolo et al., 2019). Modifications in the gut microbiome were evaluated by dosage of serum concentrations of metabolites produced by commensal bacteria such as propionate and butyrate, two major short chain fatty acids (SCFA) as well as lactate produced by lactic acid bacteria (LAB). Serum propionate, butyrate and lactate were reduced in OVX mice, and treatment with *Sb* reversed this decrease, although not reaching statistical significance for lactate and butyrate (Figure 3b-d).

Interestingly, in *Sb*-treated OVX mice we also observed a decreased proportion of BM CD4^+^ TNFα-producing T cells that have been reported to be responsible for increased osteoclastogenesis in OVX mice (Cenci et al., 2000) (Figure 3e). Lastly, we evaluated i-OCL differentiation by analyzing BM-derived OCLs for their expression of Cx3cr1, the previously described marker for i-OCLs (Ibáñez et al., 2016). In *Sb*-treated OVX mice, the proportion of Cx3cr1^+^ i-OCLs was significantly reduced compared to non-treated OVX mice, while there was no alteration of Cx3cr1^+^ OCLs in SHAM mice (Figure 3f).

### Stimulation of TLRs and CLRs influences i-OCL differentiation

Our results showed that the beneficial effect of *Sb* in OVX mice were in part due to a systemic effect on inflammatory parameters responsible for pathological osteoclastogenesis. However, yeast derivatives such as ß-glucans are known to translocate to the blood and organs through the gut barrier (Isnard et al., 2021; Rice et al., 2005), and can therefore directly affect cells expressing PRR that recognize them. Thus, we determined the effect of agonists of these receptors on the differentiation of t-OCLs and i-OCLs. We used curdlan and zymosan as Dectin-1 and TLR2 agonists and glucosyl-6-tetradecyloctadecanoate (GlcC_14_C_18,_ a synthetic C6-branched glycolipid) to stimulate Mincle. The formation of t-OCLs from BM CD11b^+^ cells was not affected by any of these agonists (Figure 4a-c), consistent with their low expression of Dectin-1, TLR2 and Mincle receptors (Figure 1j). In contrast, curdlan, zymosan and GlcC_14_C_18_ dramatically inhibited the differentiation of BM-derived DCs into i-OCLs (Figure 4a-c). To confirm the involvement of Dectin-1, TLR2 and Mincle in these effects we used neutralizing antibodies and siRNA. Anti-Dectin-1 and anti-Mincle antibodies reversed the inhibitory effect of curdlan and GlcC_14_C_18_, respectively, on i-OCL formation, compared to the control isotype (Suppl. Fig 4a-b). TLR2 siRNA was able to abrogate the decrease of i-OCL differentiation induced by curdlan, but not by zymosan (Suppl. Fig 4c). These data revealed that Dectin-1, TLR2 and Mincle activation specifically inhibits the formation of i-OCLs without affecting t-OCLs. Additionally, we performed *in vitro* assays to assess the effect of these agonists on mature i-OCL activity. Our results revealed that, in addition to blocking i-OCL differentiation, agonists of these receptors also reduced their capacity to degrade mineralized matrix (Suppl. Figure 4d).

**Figure 4:**
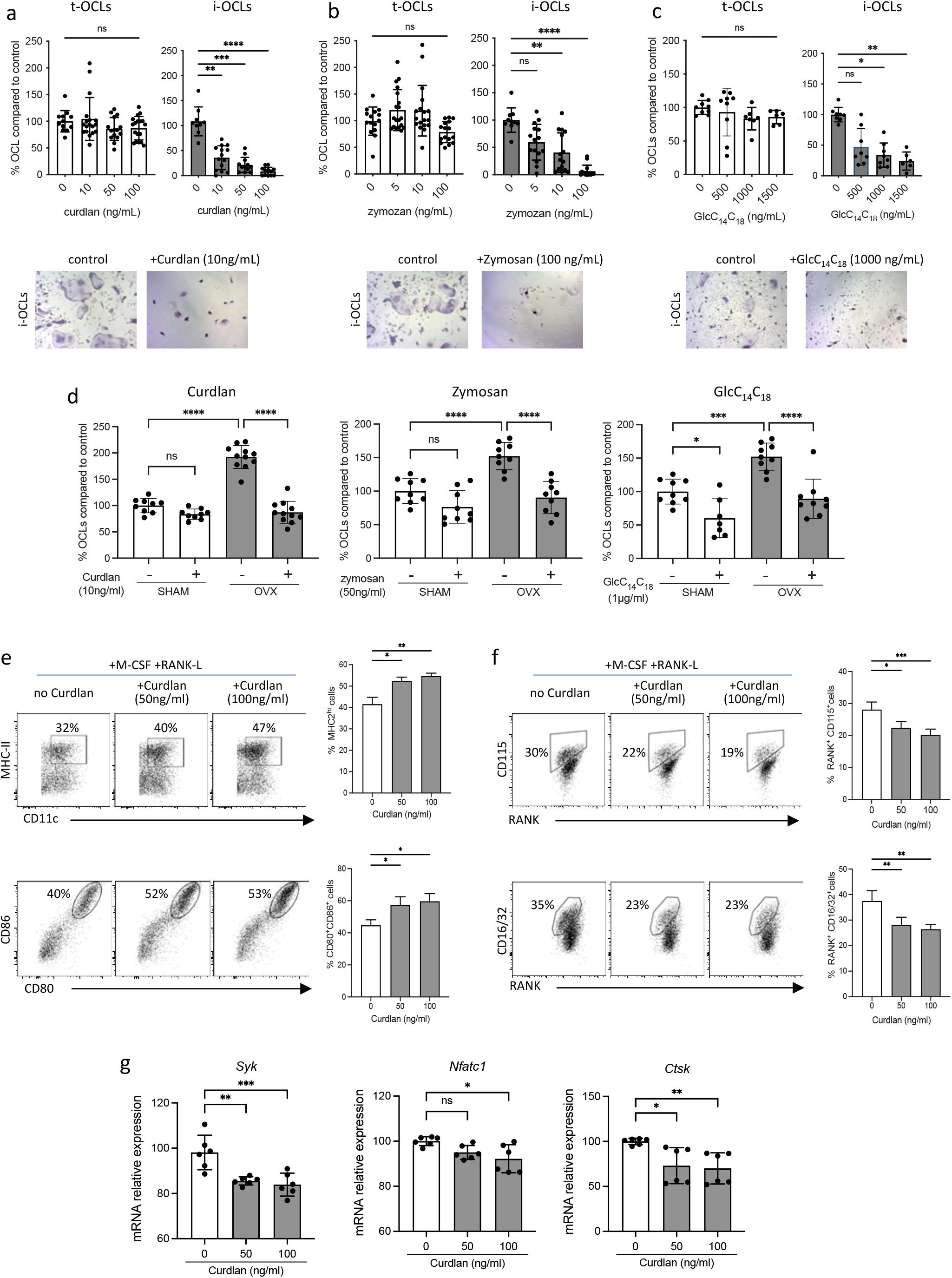
Implication of TLR2, Dectin-1 and Mincle in the differentiation of inflammatory OCLs. **(a-c)** Quantification of the differentiation of t-OCLs and i-OCLs in the presence of indicated concentrations of curdlan, zymosan and GlcC_14_C_18_. Upper panels: TRAcP^+^ cells with 3 or more nuclei were counted as OCLs. Bottom panels: representative image of TRAcP staining for the control (without agonist) and agonist-treated i-OCLs at the indicated concentration. **(d)** Enumeration of *in vitro* differentiated OCLs from BM cells of OVX and SHAM mice in the presence of indicated concentrations of curdlan, zymosan and GlcC_14_C_18_ (Glc). TRAcP^+^ cells with 3 or more nuclei were counted as OCLs (n=8-11). **(e-f)** FACS plots and quantification of the expression of **(e)** MHC-II, CD80 and CD86 and **(f)** CD115 (Csfr1), Rank and CD16/32 (FcgrII/III) on BM-DCs (n=4 biological replicates per group) cultured in osteoclast differentiation medium and stimulated or not for 24h with the indicated curdlan concentrations **(g)**RT-qPCR analysis of the expression of *Syk, Nfatc1* and *Ctsk* on BM-DCs cultured in osteoclast differentiation medium and stimulated or not for 72h with the indicated curdlan concentrations. *p<0.05; **p<0.01; ***p<0.001; ****p<0.0001; n.s. non-significant differences.

In line with the aforementioned results, the Dectin-1 and TLR2 agonists (curdlan and zymosan) also strongly reduced the differentiation of OCLs derived from BM cells of OVX mice *in vitro*, while they had no significant effect on the differentiation of OCLs generated from SHAM control mice (Figure 3d). However, GlcC_14_C_18_ reduced OCL differentiation of progenitors from OVX mice, but also to a lesser extent from SHAM mice (Figure 4d), according to the equivalent expression level of Mincle in these cells (Figure 2a).

Downstream signaling of Dectin-1 and Mincle largely involves activation of the kinase spleen tyrosine kinase Syk. Syk is also required for the differentiation and activity of OCLs (Mócsai et al., 2004). Accordingly, we found that BM-derived DCs from CD11cΔ*Syk* mice that have selective depletion of Syk in CD11c^+^ cells (Iborra et al., 2012), failed to differentiate into OCLs (Suppl. Fig 4e), indicating that Syk is required for i-OCL formation. As expected, FACS analysis showed that addition of curdlan to the OCL differentiation medium rapidly induced Syk phosphorylation (Suppl. Fig 4f), which was followed at 24h by an increased proportion of MHC-II^+^ and CD80^+^CD86^+^ DCs revealing the maturation of BM-derived DCs (Fig 4e). This treatment simultaneously reduced the proportion of Rank^+^ cells expressing Csf1r (CD115) and FcgRII/III (CD16/32), all required for OCL differentiation (Fig 4f). Furthermore, it also down regulated the expression of *Syk*, as well as its downstream targets *Nfatc1* and *Ctsk* (Fig 4g). These results show that despite a rapid activation of Syk in BM-DCs treated with M-CSF and RANK-L upon addition of curdlan, Syk expression decreases with time, as previously shown (Yamasaki et al., 2014), as well as the capacity of BM-DCs to differentiate into i-OCLs.

To investigate whether yeast probiotics have the same effect as the PRR agonists on the differentiation of i-OCLs, we used *Sb-*conditioned medium (*Sb*-CM). *Sb*-CM completely blocked the differentiation of i-OCL progenitors while it did not affect the formation of t-OCLs, except at very high concentrations (Figure 5a). This inhibitory effect was not associated with an increase in cell apoptosis (data not shown). Next, we addressed the involvement of Dectin-1, TLR2 and Mincle in the inhibitory effect of *Sb*-conditioned medium on i-OCLs as described above. While TLR2-siRNA had no effect (Figure 5b), anti-Dectin-1 and anti-Mincle blocking antibodies completely abrogated the inhibitory effect of *Sb*-CM on the differentiation of i-OCLs, demonstrating the prominent role of these receptors in mediating the effect of *Sb* on inflammatory osteoclastogenesis (Figure 5c). Moreover, as curdlan, *Sb-*CM strongly stimulated BM-DC maturation (Figure 5d) while it dramatically decreased the proportion RANK^+^ Csf1r^+^ (CD115^+^) and FcgRII/III^+^ (CD16/32^+^) cells representing OCL progenitors (Figure 5e). Lastly, *Sb* also reduced the resorption capacity of OCLs in vitro (Suppl. Fig 4d). These results revealed that derivatives from *Sb* recognized by PRRs, such as ß-glucans, interfere with the capacity of BM-derived DCs to differentiate into i-OCLs and with the activity of these OCLs.

**Figure 5:**
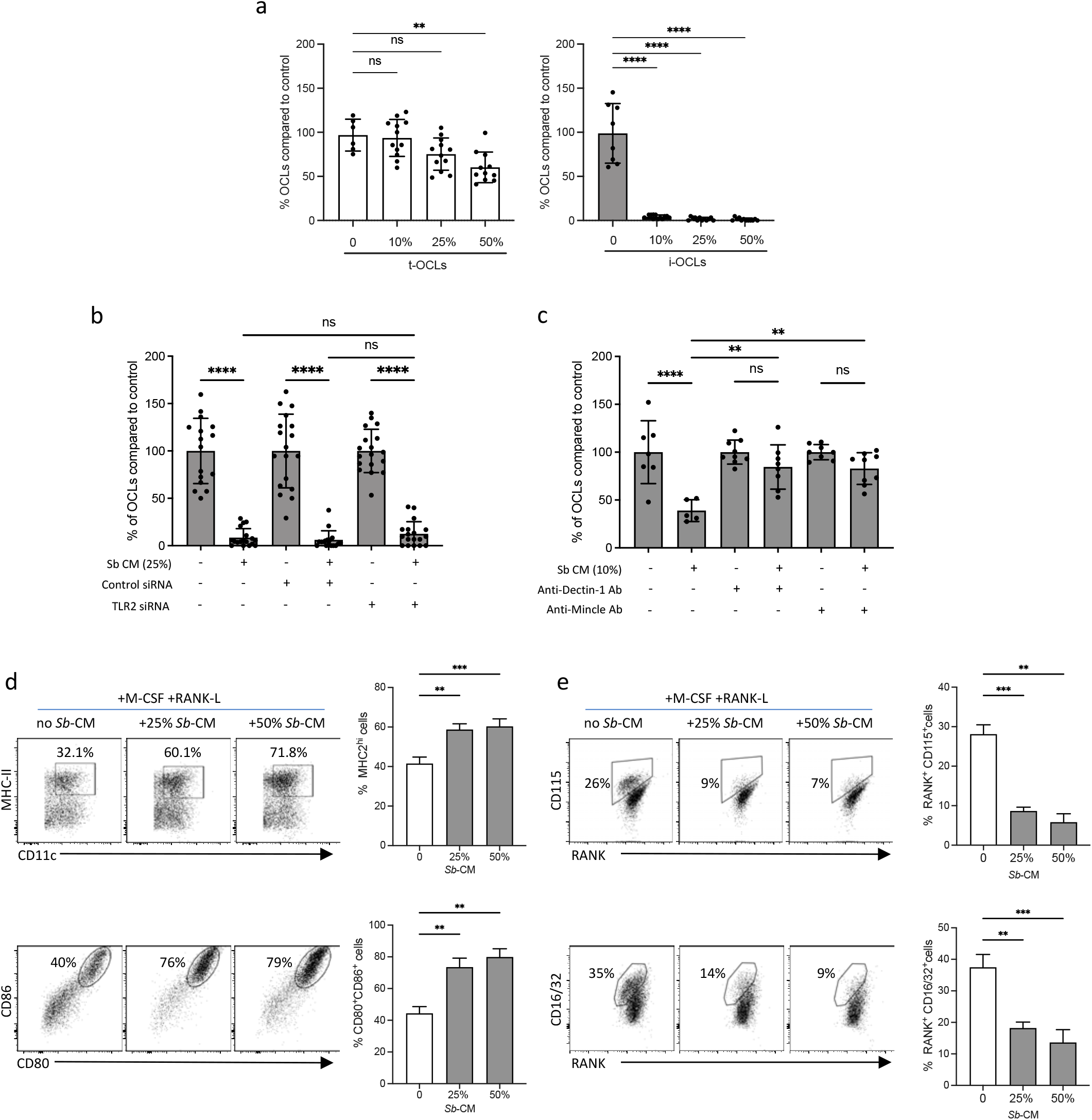
The probiotic yeast *Sb* modulates the differentiation of i-OCLs. **(a)** Quantification of differentiated t-OCLs and i-OCLs in the presence of indicated concentrations of *Sb*-conditioned medium (*Sb*-CM). **(b)** Differentiation of i-OCLs was performed in the presence of control siRNA or siRNA targeting TLR2 in the absence or presence of *Sb*-CM. (**c**) Differentiation of i-OCLs was performed in the presence of blocking antibodies for Dectin-1 and Mincle in the absence or presence of *Sb*-CM. For all experiments, TRAcP^+^ multinucleated cells with ≥3 nuclei were considered as OCLs and enumerated after 6 days of culture. **(d-e)** FACS plots and quantification of the expression of **(d)** MHC-II, CD80 and CD86 and **(e)** CD115 (Csfr1), Rank and CD16/32 (FcgrII/III) on BM-DCs (n=4 biological replicates) cultured in osteoclast differentiation medium and stimulated or not for 24h with the indicated % of *Sb*-CM. The controls panels (without *Sb*) are identical to those of Fig3.d-f as the experiments were performed together. *p<0.05; **p<0.01; ***p<0.001; ****p<0.0001; n.s. non-significant differences.

## Discussion

The present study demonstrates that i-OCLs and t-OCLs are two distinct OCL populations with specific molecular signatures that differ in their preferential use of differentiation pathways and their capacity to sense the environment through the PRRs Dectin-1, TLR2 and Mincle. Based on these specificities, we could show that targeting these receptors reduces bone loss in OVX mice and inhibits specifically the differentiation of i-OCLs while sparing t-OCLs.

Osteoclast heterogeneity has been neglected for a long time and the mechanisms regulating the emergence and function of i-OCLs are still largely unknown. Our transcriptomic profilings show that while being comparable in their expression of bone resorption-associated genes, t-OCLs and i-OCLs differ in their expression of genes associated with immune responses, which provides additional evidence for their previously reported divergent immune properties (Ibáñez et al., 2016). It also highlights differences in genes related to OCL differentiation. Osteoclastogenesis is regulated by two main pathways, the classical RANK-associated and the co-stimulatory Ig-like receptor-associated pathways, both of which are required to promote OCL differentiation (Humphrey and Nakamura, 2016; Koga et al., 2004; Seeling et al., 2013). Increased RANK-L levels as well as stimulation of FcγR and ITAM signaling are involved in pathological bone resorption in rheumatic diseases such as rheumatoid arthritis (Herman et al., 2008; Ochi et al., 2007). However, to date these pathways have not been described to be preferentially used by specific populations of OCLs. Here, we show that the implication of the co-stimulatory pathway in diseases associated with high bone resorption could be linked to its up-regulation during i-OCL differentiation, which suggests that specific targeting of this pathway and its related molecules could limit inflammatory bone loss with a minimal impact on OCLs involved in physiological bone remodeling.

Remarkably, TLR2, Dectin-1 and Mincle are increased in i-OCLs and OCLs from OVX mice compared to t-OCLs and to those from SHAM mice. These receptors are expressed in myeloid cells and sense microbial structures from bacteria, fungi or parasites, including lipoproteins, glycoproteins, peptidoglycan for TLR2, β-glucans for Dectin-1 and TLR2, and α-mannose recognized by Mincle (Sancho and Reis e Sousa, 2012; Savva and Roger, 2013). Moreover, they interact with each other and with Fcγ receptors for the induction of inflammatory responses (Gantner et al., 2003; Sato et al., 2003). Very few studies have investigated the effect of microbial β-glucans and α-mannose on osteoclastogenesis. TLR2 stimulation was shown to reduce osteoclastogenesis from non-committed OCL progenitors but to increase OCL survival and differentiation of RANK-L-primed pre-OCLs (Souza and Lerner, 2019; Takami et al., 2002). High concentrations of curdlan (in the range of μg/mL) have been reported to decrease the differentiation of OCLs from Dectin-1-transfected RAW cells or from BM cells by upregulating *MafB* and reducing *Nfatc1* expression (Zhu et al., 2017). Interestingly, here we show that the yeast probiotic *Sb* significantly reduces bone loss in OVX mice. *Sb* has proven its beneficial probiotic effects in a several of gastrointestinal disorders. It favors regeneration of the integrity of the epithelial barrier in colitis and restores the gut microbiota after antibiotic treatment or diarrhea by increasing SCFA-producing bacteria such as *Lachnospiraceae, Bacteroides, Ruminococcus* and *Prevotellaceae* (Moré and Swidsinski, 2015; Terciolo et al., 2019; Yu et al., 2017). In line with this, our results demonstrate that *Sb* improves the gut barrier function in the context of OVX-induced osteoporosis. It also normalizes the concentration of SCFAs and lactate in OVX mice, which strongly suggests that *Sb* favors bacteria species producing these metabolites. Indeed previous studies reported that the diversity of intestinal microbiota is lower in OVX mice than in SHAM mice, and its composition is altered (Li et al., 2021; Zaiss et al., 2019). Lactic acid bacteria as well as SCFA-producing bacteria have a beneficial effect on bone (Zaiss et al., 2019) which probably participates in the reduced bone loss observed in *Sb*-treated OVX mice. Moreover, gut dysbiosis and increased intestinal permeability reported in osteoporosis (He et al., 2020; Xu et al., 2020) induce activation of osteoclastogenic CD4^+^ T cells over-producing RANK-L and TNFα (D’Amelio et al., 2008; Li et al., 2016). Thus, the reduction in BM TNFα-producing CD4^+^ T cells observed after *Sb* treatment in OVX mice is likely due to its beneficial effect on the gut, as described for bacterial probiotics (Li et al., 2016). As TNFα^+^CD4^+^ T cells are major inducers of i-OCLs (Ciucci et al., 2015; Ibáñez et al., 2016; Madel et al., 2019), this effect probably also participates in the decreased proportion of i-OCLs observed in *Sb*-treated OVX mice. In addition, in various models of gut disorders, *Sb* it has been described to increase the production of anti-inflammatory cytokines such as IL-10 and IL-4 (Czerucka and Rampal, 2019; Pais et al., 2020) that are potent inhibitors of OCL formation (Fujii et al., 2012; Park-Min et al., 2009).

In addition to its systemic effects on inflammation and microbiota, *Sb* is also likely to directly affect i-OCL differentiation. The yeast cell wall components ß-glucans are well known to cross the gut barrier and translocate to the blood from where they can disseminate to organs (Isnard et al., 2021; Rice et al., 2005) and can directly modulate cells from the monocytic family (Ibáñez et al., 2019a), which includes circulating OCL progenitors, through stimulation of CLRs. Consequently, *in vivo* they can possibly exert the same direct inhibitory effect on the differentiation of i-OCLs as observed *in vitro*. This is confirmed by our in vitro analysis showing a specific inhibition of i-OCL differentiation compared to t-OCLs by *Sb*, as well as agonists of TLR2, Dectin-1 and Mincle, which is in line with the much higher expression of these receptors in BM-derived DCs than in BM MNs.

The kinase Syk plays a major role in Dectin-1 and Mincle signaling pathways and indeed, Syk is rapidly phosphorylated upon curdlan stimulation in BM-DCs. On the other hand, Syk is required for efficient OCL differentiation (Zou et al., 2007), including from DCs as shown here. However, in DCs stimulated with curdlan in osteoclastogenic medium, the expression of *Syk* decreases with time together with *Nfatc1*, a master gene of OCL differentiation, and *Ctsk*, a main marker of OCLs required for their activity, which participates in reducing i-OCL formation. These results are in agreement with the literature showing in the same conditions a degradation of Syk after treatment with high doses of curdlan (Yamasaki et al., 2014). In DCs, the interaction between TLR2 and Dectin-1 is involved in the stimulation of NF-κB and in the production of inflammatory cytokines such as TNFα and IL-12 (Gantner et al., 2003; Sancho and Reis e Sousa, 2012). This strongly induces DC activation and therefore potent phagocytic and anti-microbial responses (Gantner et al., 2003; Sancho and Reis e Sousa, 2012). Accordingly, we show that treatment with curdlan or with *Sb*-CM induces DC maturation while it decreases the proportion of Rank^+^ cells expressing Csf1r and FcgrII/III, all of which are required for OCL formation. Thus, stimulation of DCs with the PRR agonists has a dual effect, i.e. increasing their maturation and simultaneously reducing their ability to give rise to i-OCLs.

In conclusion, we here identified that i-OCLs differ from t-OCLs in the control of their differentiation and in their capacity to sense their environment and in particular to respond to stimuli through CLR and TLR activation. Based on these properties, we demonstrated that the probiotic yeast *Sb* has a beneficial effect on bone loss in osteoporotic mice by restoring gut barrier integrity and reducing osteoclastogenic CD4^+^ T cells thereby reducing indirectly i-OCLs. But it also directly targeting of i-OCL progenitors to interfere with their differentiation. These insights open new interesting perspectives for the treatment of pathological bone resorption by demonstrating that the Ig-like receptor costimulatory pathway and the associated PRR pathway, rather than the RANK pathway, could represent efficient therapeutic targets to specifically impact on inflammatory osteoclastogenesis while maintaining physiological OCL resorption. These novel insights and regulatory mechanisms mediated by yeast probiotics could provide new therapeutic options to overcome the global inhibition of OCLs and the resulting impaired bone quality associated with current anti-resorptive therapies.

## Materials and Methods

### Mice and ovariectomy-induced osteoporosis

C57BL/6 mice were purchased from Charles River Laboratory at 4-weeks of age and housed in the Animal Facility of the University Côte d’Azur. Animals were maintained under a 12h light/dark cycle and food and tap water were provided *ad libitum*. Female 6-week old C57BL/6 mice were randomly assigned into two groups for subsequent bilateral ovariectomy or SHAM surgery. Starting from two weeks after surgery, mice received by gavage *Saccharomyces boulardii* (Biocodex, Gentilly, France) 3g/kg of body weight, 3 times per week until the end of the experiment. Six weeks after surgery, mice were sacrificed. CD11cΔ*Syk* mice and CD11c-Cre littermates (Iborra et al., 2012) female mice were bred in Centro Nacional de Investigaciones Cardiovasculares (CNIC) in SPF conditions. All experiments were approved by and conducted in accordance with the Institutional Ethics Committee on Laboratory Animals (CIEPAL-Azur, Nice Sophia-Antipolis, France).

### Primary cell culture and osteoclast differentiation

Inflammatory (i-OCLs) and tolerogenic osteoclasts (t-OCLs) were differentiated *in vitro* as described previously (Halper et al., 2021; Ibáñez et al., 2016) from 6-week old C57BL/6 mice (Suppl. Fig 1a). Briefly, CD11c^+^ BM-derived DCs cells were obtained by culturing 5×10^5^ BM cells/well in 24-well plates in RPMI medium (ThermoFisher Scientific) supplemented with 5% serum (Hyclone, GE Healthcare), 1% penicillin-streptomycin (ThermoFisher Scientific), 50 μM 2-mercaptoethanol (ThermoFisher Scientific), 10 ng/ml GM-CSF and 10 ng/ml IL-4 (both PeproTech). CD11c^+^ DCs were isolated using biotinylated anti-CD11c (1:200; clone HL3; BD Biosciences) and anti-biotin microbeads (Miltenyi Biotec). Inflammatory OCLs were differentiated by seeding a total of 2×10^4^ CD11c^+^ DCs/well on 24-well plates in MEM-alpha (ThermoFisher Scientific) including 5% serum (Hyclone, GE Healthcare), 1% penicillin-streptomycin, 50 μM 2-mercaptoethanol, 25 ng/ml M-CSF and 30 ng/ml RANK-L (both R&D) (OCL differentiation medium). For t-OCL culture, 2×10^5^ CD11b^+^ monocytic BM cells that were isolated by biotinylated anti-CD11b (1:100; clone M1/70; ThermoFisher Scientific) and anti-biotin microbeads (Miltenyi Biotec), were seeded per well on 24-well plates in OCL differentiation medium as described above. For OCL differentiation from OVX and SHAM control mice, 5×10^5^ BM cells were cultured per well in 24-well plates in OCL differentiation medium as indicated above. When indicated, differentiation was performed in the presence of indicated concentrations of curdlan, zymosan or GlcC_14_C_18_. Medium conditioned by *Saccharomyces boulardii* CNCM I-745 was prepared by culture of the yeast in MEM-alpha overnight. The medium was collected and filtered through a sterile 22μm filter, complemented with serum and antibiotics as described above and used as conditioned medium at indicated dilutions. OCL differentiation was evaluated at the end of the differentiation after TRAcP staining according to manufacturer’s instructions (Sigma-Aldrich). Mature OCLs were enumerated under a light microscope as multinucleated (≤3 nuclei/cell) TRAcP^+^ cells. OCL activity was evaluated by seeding OCL progenitors on plates coated with resorbable matrix (Osteoassay) in OCL differentiation medium. Resorbed areas were quantified (ImageJ software version 1.53, NIH, Bethesda, MD) after removing of the cells with water and staining of the mineralized matrix with alizarine red.

### RNA-sequencing

RNAseq analysis was performed on mature multinucleated OCLs. After differentiation *in vitro*, t-OCLs and i-OCLs (5 biological replicates, each derived from one mouse) were detached with Accutase, labeled with 5 μg/ml H33342 and sorted on their multinucleation as previously described (Madel et al., 2018) (c.f. Suppl Figure 1b for gating strategy). Total RNA (100 ng) was extracted from sorted OCLs (RNeasy kit, Qiagen) and directional libraries were prepared (Truseq stranded total RNA library kit, Illumina). Libraries were pooled and sequenced paired-ended for 2×75 cycles (Nextseq500 sequencer, Illumina). 30-40 million fragments were generated per sample and quality controls were performed. Data were analyzed by 2 approaches as described previously (Madel et al., 2020). Both gave equivalent results. For the first one, reads were “quasi” mapped on the reference mouse transcriptome (Gencode vM15) and quantified (SALMON software, mapping mode and standard settings) (Patro et al., 2017). Transcript count estimates and confidence intervals were computed using 1.000 bootstraps to assess technical variance. Transcript counts were aggregated for each gene for computing gene expression levels. Gene expression in biological replicates (n=5) was then compared between t-OCLs and i-OCLs with a linear model (Sleuth (Pimentel et al., 2017)) and a false discovery rate of 0.01. For the second approach, raw FASTQ reads were trimmed with Trimmomatic and aligned to the reference mouse transcriptome (Gencode mm10) with STAR (Dobin et al., 2013) on the National Institutes of Health high-performance computing Biowulf cluster. Gene-assignment and estimated read counts were assessed using HTseq (Anders et al., 2015). Gene expression was compared between t-OCLs and i-OCLs using DESeq2 (Love et al., 2014) with the Wald test (FDR < 0.01). Lists of differentially expressed genes were annotated using Innate-DB and EnrichR web portals.

### miRNA Profiling and validation by RT-qPCR

Total RNA from sorted OCLs was extracted using the miRNeasy Micro Kit (QIAGEN) and the procedure automatized using the QIAcube (QIAGEN). The miRNA expression profiles were analyzed on paired samples using the TaqMan® Array rodent MicroRNA Card Set v3.0 (TLDA, Applied Biosystems) after pre-amplification steps, according to manufacturer’s instructions. Relative expression and statistical analysis were calculated using the

ExpressionSuite software (Applied Biosciences), which included the student’s t-test for sample group comparisons and built Volcano Plot comparing the size of the fold change (biological significance) to the statistical significance (p-value). Dysregulated miRNAs were examined with MirWalk (Dweep and Gretz, 2015), a miRNA database aiming to identify predicted and validated target genes and related pathways. This software provides information on miRNA-target interactions, not only on 3′-UTR, but also on the other regions of all known genes, and simultaneously interrogates several algorithms (TargetScan, Miranda, RNA22 and miRWalk). We used a high predictive score with at least 3 of the 4 queried algorithms predicting miRNA target genes. Comparison of the expression patterns of 750 miRNAs in i-OCLs and t-OCLs was performed using Ct values <35, difference of at least 2-fold with a p-value <0.05.

Mature miRNAs of interest were specifically converted into cDNA using TaqMan microRNA reverse transcription kit according to the manufacturer’s protocol (Applied Biosystems). RT specific primers 5X (ThermoFisher) were multiplexed in a primer pool containing 1% of each diluted in an adequate volume of Tris-EDTA 1X. Pre-amplification step was performed using FAM-labeled specific PCR primers 20X and TaqMan PreAmp Master Mix kit (Applied Biosystems) for 12 cycles. Alternatively, these preliminary steps were performed using Megaplex RT and PreAmp Primers, Rodent pool A (Applied Biosystems) that include specific primers for miRNAs of interest. Quantitative Real Time PCR (RT-qPCR) was performed on diluted pre-amp products using the specific TaqMan PCR primers and TaqMan Universal Master Mix II with no UNG, and run on Viia7 system (Applied Biosystems) in 96-well PCR plates for 40 cycles. Relative miRNA expression was normalized on sno202 expression in murine cells with the 2^-ΔCT^ method.

### Bone structure analyses

Long bones of OVX and SHAM-operated mice were fixed in 4% paraformaldehyde. Bone microarchitecture analysis using high-resolution microcomputed tomography (μCT) was performed at the pre-clinical platform ECELLFRANCE (IRMB, Montpellier, France). Cortical and trabecular femora were imaged using high-resolution μCT with a fixed isotropic voxel size of 9 μm with X-ray energy of 50 kV, current of 500 mA, 0.5 mm aluminum filter and 210 ms exposure time. For visual representation, 3D reconstructions were generated using NRecon software (Bruker μCT, Belgium).

### FITC-Dextran permeability assay

One hour before sacrifice, mice received oral gavage of 3-5 kDa fluorescein isothiocyanate (FITC)–dextran (Sigma-Aldrich) (60 mg/100 g body weight). FITC-dextran concentration in serum was measured by fluorometry in a fluorimeter (Xenius, SAFAS, Monaco) at 488/525 nm. Standard curve was prepared using dilutions of FITC-dextran in PBS with 20% fetal calf serum.

### TRAcP staining and Immunohistochemistry

Tibias were fixed in 4% paraformaldehyde and decalcified in 4.13% EDTA, 0.2% paraformaldehyde pH 7.4, at 50°C in KOS microwave tissue processor (Milestone, Michigan, USA). They were then dehydrated and embedded in paraffin. TRAcP staining was performed as described (Lézot et al., 2015) with Mayer hematoxylin counterstaining on 3 μm thick sections to identify osteoclasts. Immunostaining of osteoblasts and osteocytes were performed with rabbit polyclonal anti-Osterix antibody (ab22552, dilution 1/800 ; Abcam, Cambridge, UK) and rabbit polyclonal anti-sclerostin antibody (AF1589, dilution1:200 ; R&D System, Abingdon, UK), respectively, with Gill2 hematoxylin counterstaining. Stained sections were automatically numerized (nanozoomer, Hamamatsu photonics) before observation (NDP view virtual microscope, Hamamatsu) and quantification (ImageJ software version 1.53, NIH, Bethesda, MD).

### Dosage of biochemical parameters

Concentration of lactate, propionate and butyrate in the serum was determined by ion chromatography analysis after depletion of proteins and lipids with acetonitrile (Sigma-Aldrich). Samples were loaded on a Dionex ICS-5000 Plus system automatic device (ThermoScientific) and elution was performed according to the manufacturer’s protocol. Chromatograms were aligned to standard solutions of each compound individually. Compound concentrations were determined using Chromeleon software (Thermo Scientific) by measuring surface area under the curve of the peaks and were compared to the corresponding ion standard profiles.

Seric crosslaps (CTX) were evaluated by enzyme-linked immunosorbent assay according to the manufacturer’s protocol (RatLaps™ (CTX-I) EIA, Immunodiagnostic Systems Limited).

**Flow cytometry analysis**

CD11b^+^ monocytic BM cells and BM-derived CD11c^+^ DCs cells were analyzed for their expression of Dectin-1 (1:100; clone bg1fpj; ThermoFisher Scientific), Dectin-2 (1:100, clone 17611; R&D Systems), TLR2 (1:200; clone 6C2; BD Biosciences) and Mincle (1:50, clone 6G5, Invivogen).

For analysis of BM-derived DC maturation, BM-DCs were treated or not with curdlan or *Sb*-CM at the indicated dose for 24h in OCL differentiation medium (containing 25 ng/ml M-CSF and 30 ng/ml RANK-L). They were then labelled with anti-CD11c (1:200; Clone N418, eBioscience), MHC-II/IAb (1:200; Clone AF6-120.1, BD Bioscience), CD80 (1:100; Clone 16-

10A1, eBioscience), CD86 (1:100; Clone GL1, BD Bioscience), CD115 (1:100; Clone AFS98, eBioscience) and RANK/CD265 (1:100; clone R12-31, eBioscience). Cells were analyzed by flow cytometry (BD FACSCanto-II, BD Bioscience).

To investigate Syk phosphorylation after curdlan stimulation, BM-derived DCs were stimulated for 15 minutes in OCL differentiation medium with the indicated doses of curdlan and subsequently fixed with 2% PFA (Transcription Factor Fixation/Permeabilization kit, eBioscience) over night at 4°C. Surface staining was then performed with anti-MHC-II/IAb (1:100;Clone AF6-120.1, BD Bioscience) and CD11c (1:200; Clone N418, eBioscience) antibodies before the cells were permeabilized with 1x Saponine for 15 minutes and stained with Phospho-Syk (Tyr348, clone moch1ct, eBioscience) for 45 minutes. Cells were washed and acquired on a BD FACSCanto II.

For FACS analysis on OCLs, mature OCLs were detached using Accutase (ThermoFisher Scientific), labeled with 5 μg/ml H33342 and with anti-Dectin-1, Dectin-2, TLR2 and Mincle antibodies and analyzed after doublet exclusion as multinucleated cells with 3 or more nuclei as previously described (Madel et al., 2018) for their expression of these markers (see Suppl. Figure 1b for the gating strategy). Cells were analyzed by flow cytometry (BD FACSCanto-II, BD Bioscience).

For intracellular cytokine analysis, T cells isolated from the BM of SHAM and OVX mice were stimulated with PMA, ionomycin and brefeldin A, labeled with anti-CD4 antibody (1:1000; clone RM4-5; BD Biosciences) and fixed with 4% formaldehyde overnight as described (Ciucci et al., 2015). Cells were subsequently stained with anti-TNFα antibody (1:400; clone MP6-XT22; ThermoFisher Scientific) in saponin 1X. Data were acquired using a FACSCanto-II (BD Biosciences). All FACS data were analyzed with FlowJo 10.8.1.

### In vitro mineralization assay

Osteoblastic differentiation of BM-MSC cells from C57Bl/6 mice was performed for 3 weeks in MEM-alpha medium with 10% FBS, 1% penicillin-streptomycin, 50 μM 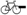mercaptoethanol supplemented with 170 μM L-ascorbic acid, 10 mM 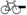glycerophosphate and 0.1 μM dexamethasone (Merck). This differentiation medium (with adjusted concentration of inductors) was supplemented with the indicated % of *Sb* conditioned medium. Medium was changed every 4 days. At day 21, cultures were fixed with 4% PFA and stained with 2% Alizarin red S (Merck). For the quantification, the Alizarin staining was dissolved in acetic acid (10%), heated 10 min at 85°C, centrifuged and buffered with ammonium hydroxide. Absorbance at 405 nm was measured (Xenius SAFAS, Monaco).

### Blocking of Dectin-1, TLR2 and Mincle

For inhibition with siRNA, BM-derived DCs were transfected with ON-TARGET SMART pool of 4 siRNA to mouse Tlr2 (50 nM, Dharmacon, Horizon Discovery, USA) using lipofectamine RNAiMAX (Invitrogen), and were further differentiated into OCLs for 5-6 days either in OCL differentiation medium in the presence of the indicated concentration of agonists, or in *Sb*-conditioned medium as described above.

For inhibition with blocking antibodies, BM-derived DCs were cultured either in OCL differentiation medium containing the indicated concentration of agonists, or in *Sb*-conditioned medium. Anti-Dectin-1 (clone bg1fpj; ThermoFisher Scientific), anti-Mincle (clone 6G5, Invivogen) or control isotype antibodies were added at the indicated concentration at the beginning of the differentiation. Medium was changed at day 3.

For both approaches, OCLs differentiation was evaluated after TRAcP staining and mature OCLs were enumerated under a light microscope as multinucleated TRAcP^+^ cells.

### RT-qPCR

BM-derived DCs were stimulated for 72 hours with the indicated doses of curdlan. Total RNA was extracted with Trizol according to the manufacturer protocol. RNAs were reverse transcribed (Superscript II, Life Technologies) and RT-PCR was performed using SYBR green and the primers indicated below. Results were normalized to the 36B4 gene with the 2^−ΔCt^ method. *Syk*: AACGTGCTTCTGGTCACACA and AGAACGCTTCCCACATCAGG; *Ctsk*: CAGCAGAGGTGTGTACTATG and GCGTTGTTCTTATTCCGAGC; *Nfatc1*: TGAGGCTGGTCTTCCGAGTT and CGCTGGGAACACTCGATAGG; *36B4*: TCCAGGCTTTGGGCATCA and CTTTATCAGCTGCACATCACTCAGA.

### Statistical analysis

Data were analyzed and statistics prepared using Graph Pad Prism 9.2 software. Analyses were done using two-tailed unpaired t-test when comparing two groups and ANOVA with multiple comparison test when comparison of more than two groups. Statistical significance was considered at p<0.05 and experimental values are presented as mean ± S.D. Biological replicates were obtained from different mice.

## Data availability

The datasets generated during the current study are available from the corresponding author on reasonable request.

## Acknowledgements

The authors would like to acknowledge the Genomic Facility of the UFR Simone Veil, Université Versailles-Saint-Quentin (France) for the RNA sequencing, the IRCAN animal core facility that is supported by “la Région Provence Alpes-Côte d’Azur” (Nice, France) as well as the Montpellier preclinical platform of ECELLFRANCE for μCT analysis (IRMB, Montpellier, France). They also thank the Laboratory of Biochemistry-Hormonology (CHU, Nice, France) for CTX dosage and M. Salah and D. Carro (LP2M, Nice France) for their technical contribution. This work utilized the computational resources of the NIH-HPC-Biowulf cluster (http://hpc.nih.gov).

The work was supported by the Agence Nationale de la Recherche (ANR-16-CE14-0030) as well as by the French government, managed by the ANR as part of the Investissement d’Avenir UCA^JEDI^ project (ANR-15-IDEX-01) and Biocodex (Gentilly, France). M-B. M. was supported by the Fondation pour la Recherche Médicale (FRM, ECO20160736019), and by awards from the Fondation Arthritis, the Société Française de Biologie des Tissus Minéralisés (SFBTM), the European Calcified Tissue Society (ECTS) and the American Society of Bone and Mineral Research (ASBMR). J.H. was supported by an award from the European Calcified Tissue Society (ECTS). T.C. was supported by the Intramural Research Program of the National Cancer Institute, Center for Cancer Research, National Institutes of Health.

## Author Contributions

C.B-W. and A.W. conceived the project. M-B.M., J.H., L.I., M.R. and C.L. performed experiments, analyzed and interpreted data. A.B. and M.T. performed *in vivo* experiments and analyzed data. D.H. supervised and J.A. performed histological analysis on osteoclasts, osteoblasts and osteocytes. D.S. and S.I. provided the CD11cΔ*Syk* mutant mice, participated in experiments on these mice and commented on the manuscript. D.C. and R.P-B participated in experiment on yeast-conditioned medium. H-J.G. supervised and C.H. performed RNA-sequencing. F.A, I.D-R and C.L performed miRNome analysis and interpreted the data. M-B.M., C.B-W, T.C., I.D-R. and H-J.G. analyzed the RNAseq data. F.A., I.D-R., D.C. and M.R. contributed to the discussion of results and provided helpful advices throughout the study. M-B.M. and C.B-W. wrote the manuscript. All authors commented on the paper.

## Supplemental data

**Supplemental Figure 1.**
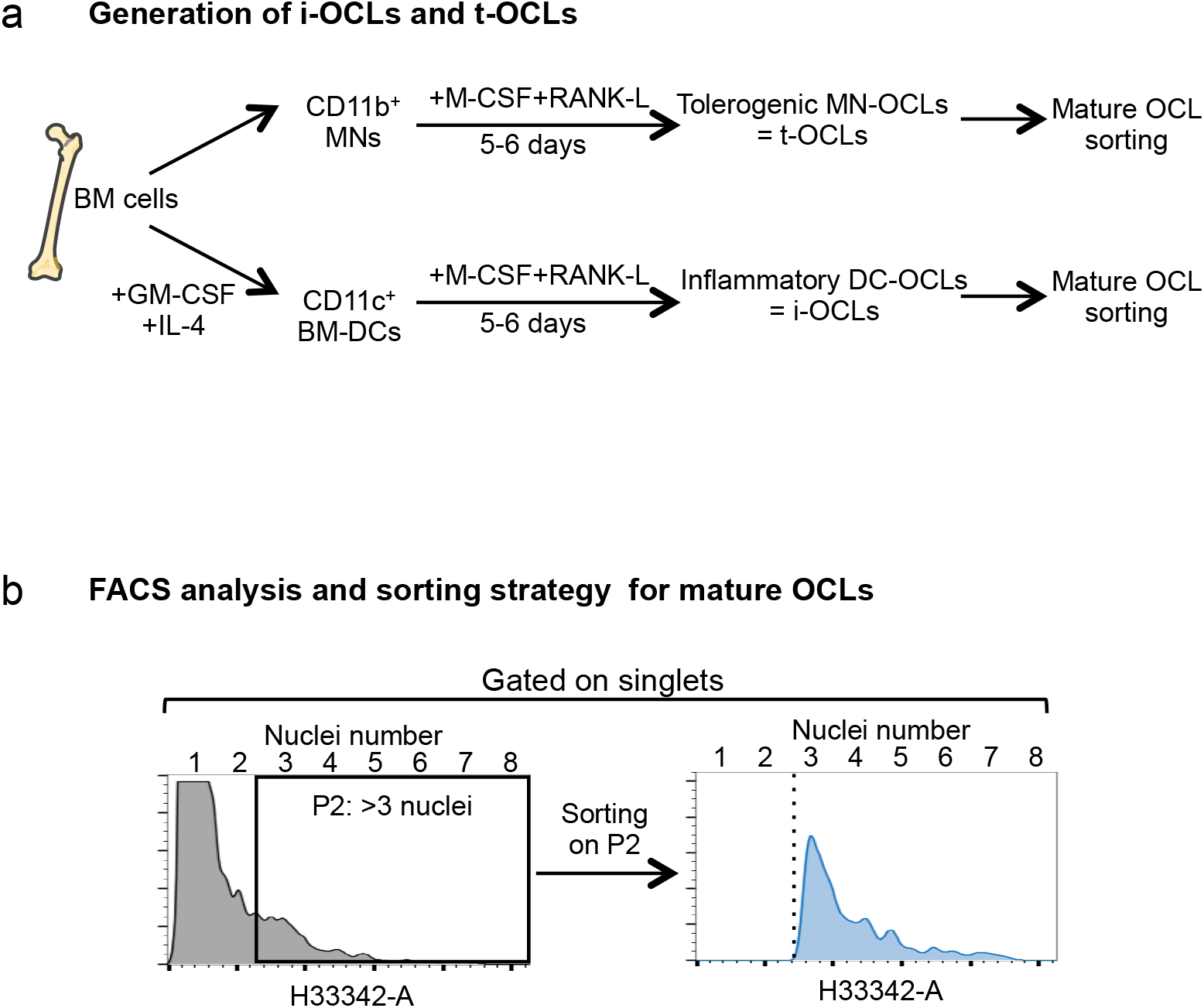
**(a)** Schematic representation of the experimental procedure for the generation of t-OCLs from BM CD11b^+^ monocytic cells (MNs) and i-OCLs from BM-derived CD11c^+^ dendritic cells (BM-DCs), as previously described (4–6). **(b)** Gating strategy for purification by cell sorting or for FACS analysis of mature osteoclasts (with 3 and more nuclei), after staining of DNA with H33342 and doublet exclusion according to the protocol we previously established (2). This strategy was used for gating on multinucleated mature osteoclasts throughout the different analyses.

**Supplemental Figure 2.**
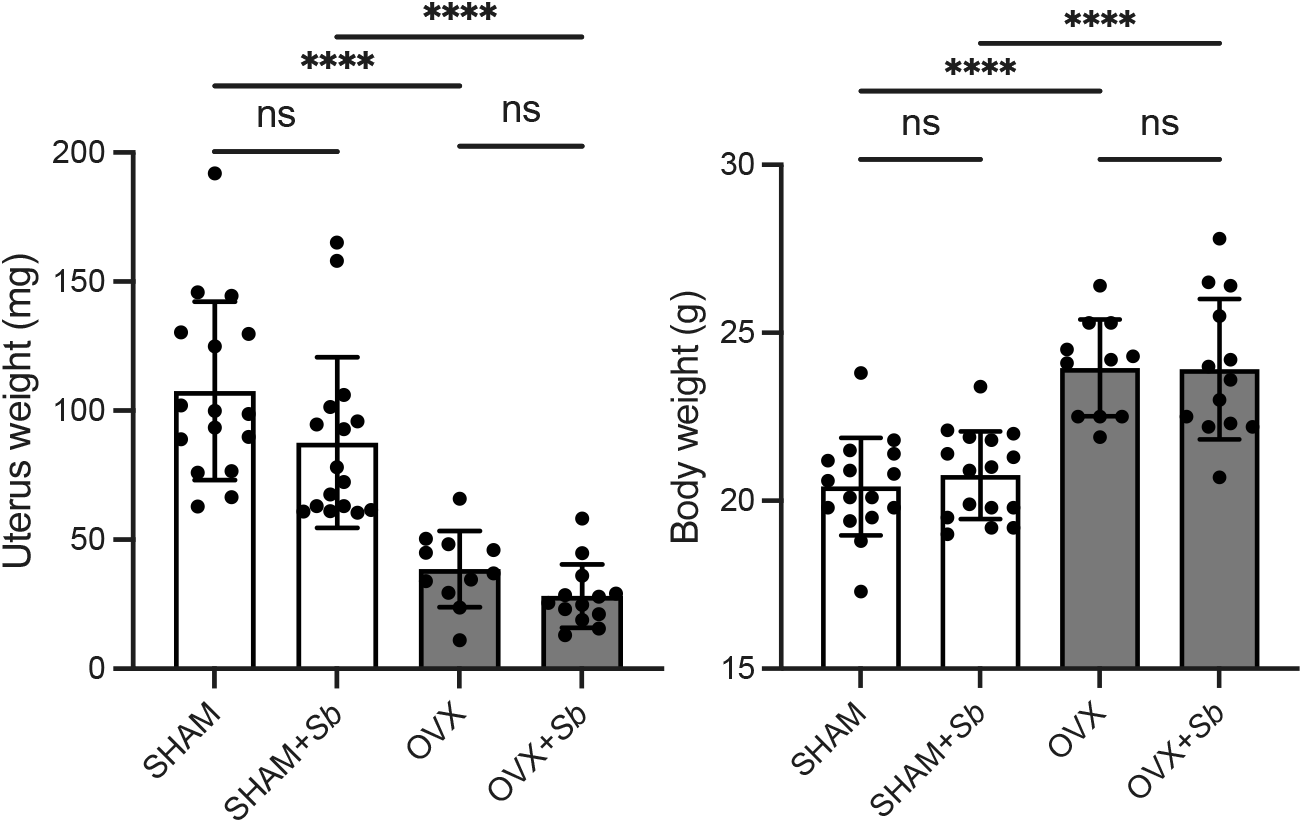
**(a)** Weight of the uteri of OVX and SHAM mice treated or not with *Sb* or *Sc*. **(b)** Body weight of the mice in the different groups, measured the day of sacrifice. Data show no differences between yeast-treated mice and control mice in each group (SHAM and OVX).). **** p<0.0001; n.s. non-significant differences.

**Supplemental Figure 3.**
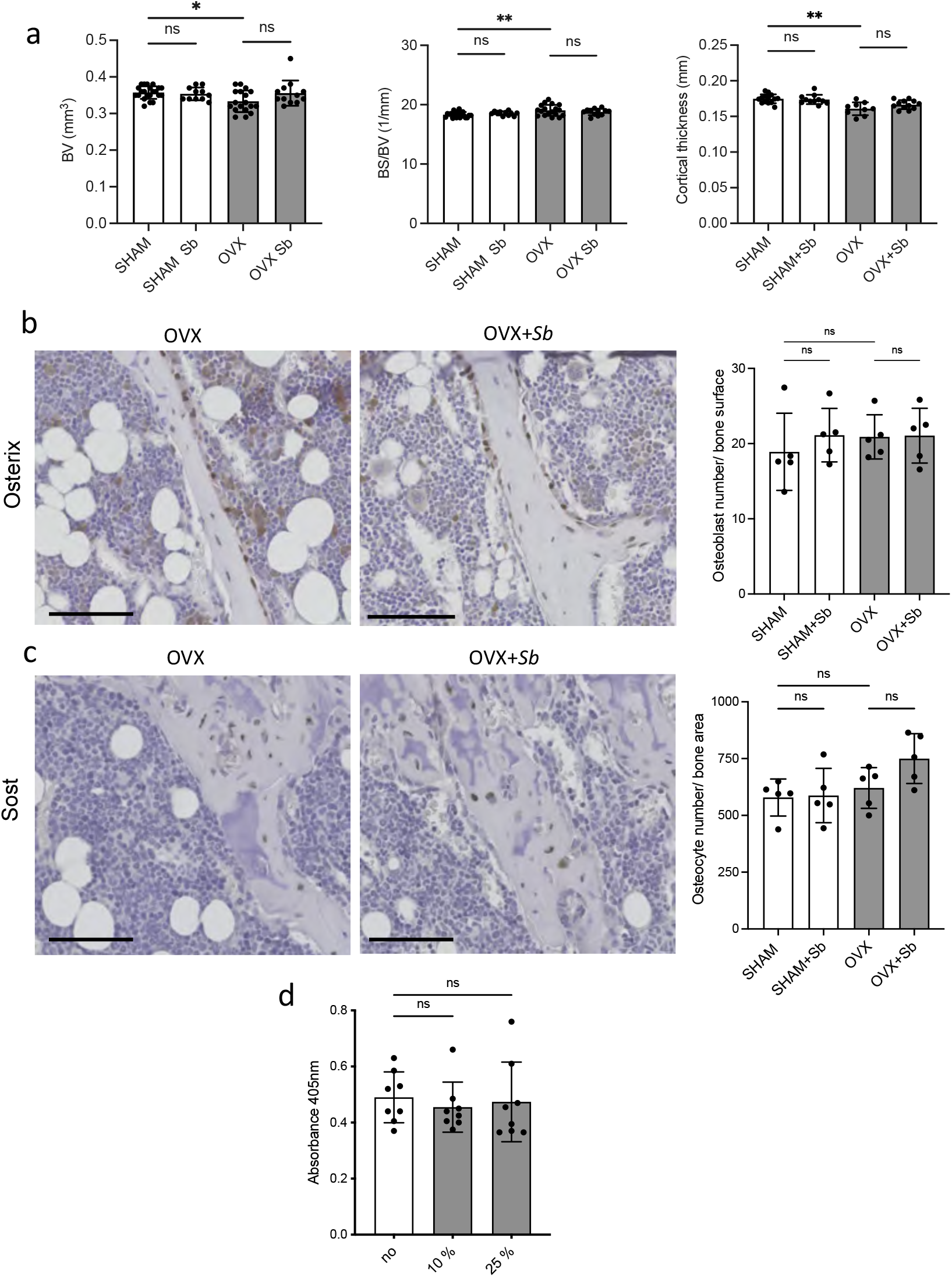
(**a**) Histograms indicate mean ± S.D. of cortical bone volume (BV), cortical thickness and cortical bone surface/bone volume (BS/BV). (**b-c**) Histological analysis of the tibia of SHAM and OVX mice treated or not with *Sb*. (**b**) Osteoblasts were identified as Osterix-expressing cells attached to the bone surface and (**c**) osteocytes were identified by their expression of Sost. Histograms represent the enumeration of the cells (n=5 mice per group). Scale bars: 100μm (**d**) *In vitro* mineralization capacity as measured by alizarin coloration of mineralized nodules deposited *in vitro* by osteoblasts treated or not with the indicated proportion of medium conditionned by *Sb* (*Sb*-CM). *p<0.05; **p<0.01; n.s., non-significant differences.

**Supplemental Figure 4.**
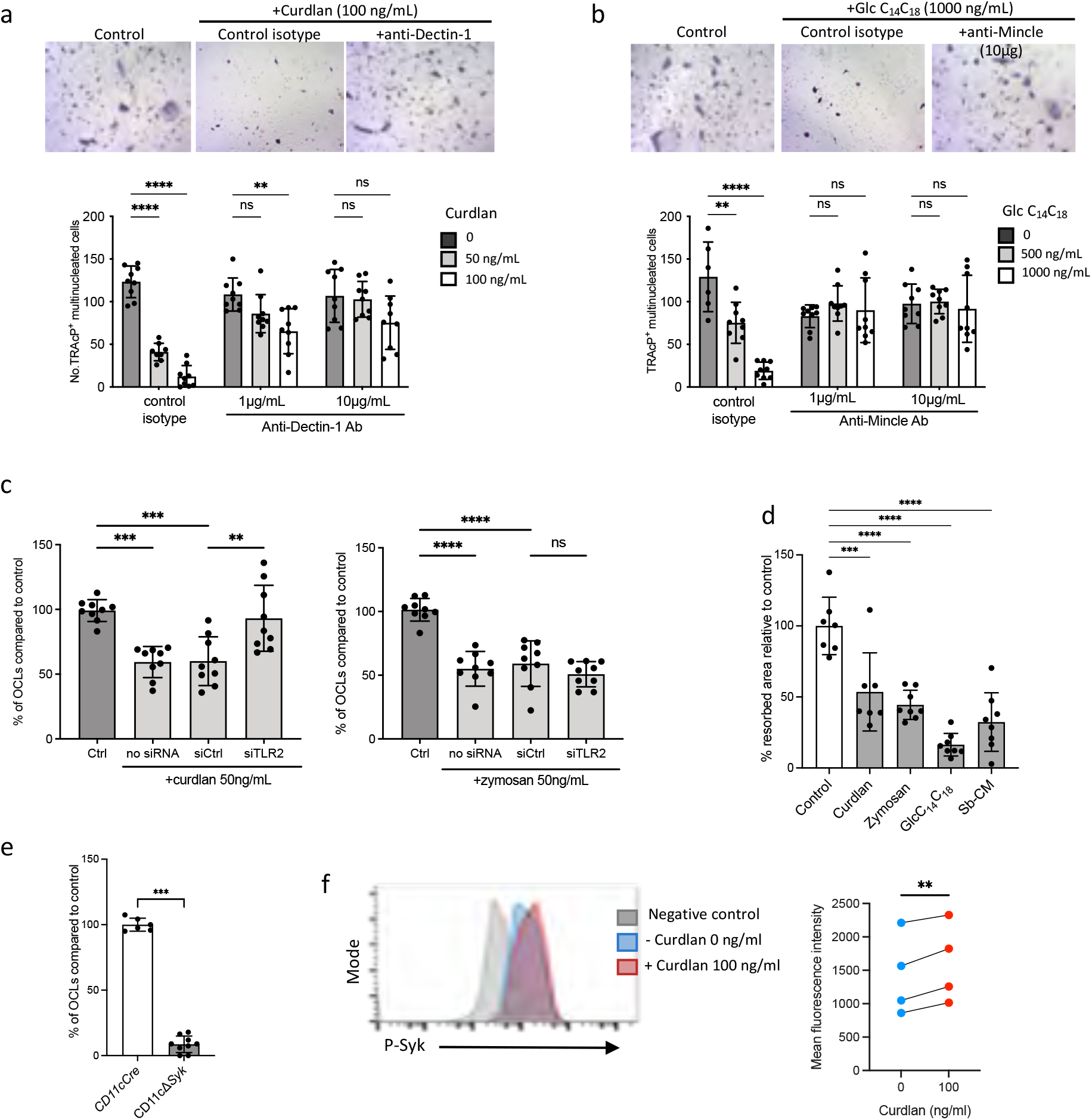
**(a-b)** TRAcP staining and quantification of OCLs (TRAcP^+^ cells ≥3 nuclei) differentiated in the presence of control isotypes or blocking antibodies against (**a**) Dectin-1 and (**b**) Mincle. **(c)** Quantification of i-OCLs differentiated in the presence of control siRNA or siRNA targeting TLR2. **(d)** Quantification of differentiation of i-OCLs from CD11cΔ*Syk* and control mice. **(e)** Flow cytometry analysis of Syk phosphorylation after 15 min of stimulation with 100ng/ml of curdlan on BM-derived DCs cultured. The right panel shows the increase in the mean fluorescence intensity (MFI) revealing increased Syk phophorylation (n=4 biological replicates). (**f**) OCL activity was evaluated by seeding BM-derived DCs on plates coated with resorbable matrix. After i-OCLs started to fuse (day 3 of differentiation), the different agonists were added at the following concentration: curdlan: 50ng/ml; zymosan, 50ng/ml, GlcC_14_C_18_: 1000ng/ml, *Sb* conditioned medium (*Sb*-CM): 25%. Resorbed area were quantified after 3 days. *p<0.05; **p<0.01; ***p<0.001; **** p<0.0001; n.s., non-significant differences.

## References

Anders S, Pyl PT, Huber W. 2015. HTSeq--a Python framework to work with high-throughput sequencing data. Bioinformatics 31:166–169. doi:10.1093/bioinformatics/btu638

Britton RA, Irwin R, Quach D, Schaefer L, Zhang J, Lee T, Parameswaran N, McCabe LR. 2014. Probiotic L. reuteri Treatment Prevents Bone Loss in a Menopausal Ovariectomized Mouse Model: PROBIOTICS SUPPRESS ESTROGEN DEFICIENCY-INDUCED BONE LOSS. Journal of Cellular Physiology 229:1822–1830. doi:10.1002/jcp.24636

Cenci S, Weitzmann MN, Roggia C, Namba N, Novack D, Woodring J, Pacifici R. 2000. Estrogen deficiency induces bone loss by enhancing T-cell production of TNF-α. Journal of Clinical Investigation 106:1229–1237. doi:10.1172/JCI11066

Ciucci T, Ibáñez L, Boucoiran A, Birgy-Barelli E, Pène J, Abou-Ezzi G, Arab N, Rouleau M, Hébuterne X, Yssel H, Blin-Wakkach C, Wakkach A. 2015. Bone marrow Th17 TNFα cells induce osteoclast differentiation, and link bone destruction to IBD. Gut 64:1072–1081. doi:10.1136/gutjnl-2014-306947

Czerucka D, Rampal P. 2019. Diversity of Saccharomyces boulardii CNCM I-745 mechanisms of action against intestinal infections. World J Gastroenterol 25:2188–2203. doi:10.3748/wjg.v25.i18.2188

D’Amelio P, Grimaldi A, Di Bella S, Brianza SZM, Cristofaro MA, Tamone C, Giribaldi G, Ulliers D, Pescarmona GP, Isaia G. 2008. Estrogen deficiency increases osteoclastogenesis up-regulating T cells activity: a key mechanism in osteoporosis. Bone 43:92–100. doi:10.1016/j.bone.2008.02.017

Dobin A, Davis CA, Schlesinger F, Drenkow J, Zaleski C, Jha S, Batut P, Chaisson M, Gingeras TR. 2013. STAR: ultrafast universal RNA-seq aligner. Bioinformatics 29:15–21. doi:10.1093/bioinformatics/bts635

Dweep H, Gretz N. 2015. miRWalk2.0: a comprehensive atlas of microRNA-target interactions. Nature Methods 12:697–697. doi:10.1038/nmeth.3485

Fujii T, Kitaura H, Kimura K, Hakami ZW, Takano-Yamamoto T. 2012. IL-4 inhibits TNF-α-mediated osteoclast formation by inhibition of RANKL expression in TNF-α-activated stromal cells and direct inhibition of TNF-α-activated osteoclast precursors via a T-cell-independent mechanism in vivo. Bone 51:771–780. doi:10.1016/j.bone.2012.06.024

Gantner BN, Simmons RM, Canavera SJ, Akira S, Underhill DM. 2003. Collaborative Induction of Inflammatory Responses by Dectin-1 and Toll-like Receptor 2. J Exp Med 197:1107–1117. doi:10.1084/jem.20021787

Halper J, Madel M-B, Blin-Wakkach C. 2021. Differentiation and Phenotyping of Murine Osteoclasts from Bone Marrow Progenitors, Monocytes, and Dendritic Cells. Methods Mol Biol 2308:21–34. doi:10.1007/978-1-0716-1425-9_2

He J, Xu S, Zhang B, Xiao C, Chen Z, Si F, Fu J, Lin X, Zheng G, Yu G, Chen J. 2020. Gut microbiota and metabolite alterations associated with reduced bone mineral density or bone metabolic indexes in postmenopausal osteoporosis. Aging (Albany NY) 12:8583–8604. doi:10.18632/aging.103168

Herman S, Müller RB, Krönke G, Zwerina J, Redlich K, Hueber AJ, Gelse H, Neumann E, Müller-Ladner U, Schett G. 2008. Induction of osteoclast-associated receptor, a key osteoclast costimulation molecule, in rheumatoid arthritis. Arthritis Rheum 58:3041–3050. doi:10.1002/art.23943

Humphrey MB, Nakamura MC. 2016. A Comprehensive Review of Immunoreceptor Regulation of Osteoclasts. Clinical Reviews in Allergy & Immunology 51:48–58. doi:10.1007/s12016-015-8521-8

Ibáñez L, Abou-Ezzi G, Ciucci T, Amiot V, Belaïd N, Obino D, Mansour A, Rouleau M, Wakkach A, Blin-Wakkach C. 2016. Inflammatory osteoclasts prime TNFα-producing CD4(+) T cells and express CX3 CR1. J Bone Miner Res. doi:10.1002/jbmr.2868

Ibáñez L, Pontier-Bres R, Larbret F, Rekima A, Verhasselt V, Blin-Wakkach C, Czerucka D. 2019a. Saccharomyces boulardii Strain CNCM I-745 Modifies the Mononuclear Phagocytes Response in the Small Intestine of Mice Following Salmonella Typhimurium Infection. Front Immunol 10:643. doi:10.3389/fimmu.2019.00643

Ibáñez L, Rouleau M, Wakkach A, Blin-Wakkach C. 2019b. Gut microbiome and bone. Joint Bone Spine 86:43–47. doi:10.1016/j.jbspin.2018.02.008

Iborra S, Izquierdo HM, Martínez-López M, Blanco-Menéndez N, Sousa CR e, Sancho D. 2012. The DC receptor DNGR-1 mediates cross-priming of CTLs during vaccinia virus infection in mice. J Clin Invest 122:1628–1643. doi:10.1172/JCI60660

Isnard S, Lin J, Bu S, Fombuena B, Royston L, Routy J-P. 2021. Gut Leakage of Fungal-Related Products: Turning Up the Heat for HIV Infection. Front Immunol 12:656414. doi:10.3389/fimmu.2021.656414

Jacome-Galarza CE, Percin GI, Muller JT, Mass E, Lazarov T, Eitler J, Rauner M, Yadav VK, Crozet L, Bohm M, Loyher P-L, Karsenty G, Waskow C, Geissmann F. 2019. Developmental origin, functional maintenance and genetic rescue of osteoclasts. Nature 568:541–545. doi:10.1038/s41586-019-1105-7

Jones RM, Mulle JG, Pacifici R. 2017. Osteomicrobiology: The influence of gut microbiota on bone in health and disease. Bone. doi:10.1016/j.bone.2017.04.009

Kiesel JR, Buchwald ZS, Aurora R. 2009. Cross-presentation by osteoclasts induces FoxP3 in CD8+ T cells. J Immunol 182:5477–5487. doi:10.4049/jimmunol.0803897

Koga T, Inui M, Inoue K, Kim S, Suematsu A, Kobayashi E, Iwata T, Ohnishi H, Matozaki T, Kodama T, Taniguchi T, Takayanagi H, Takai T. 2004. Costimulatory signals mediated by the ITAM motif cooperate with RANKL for bone homeostasis. Nature 428:758–763. doi:10.1038/nature02444

Lézot F, Chesneau J, Navet B, Gobin B, Amiaud J, Choi Y, Yagita H, Castaneda B, Berdal A, Mueller CG, Rédini F, Heymann D. 2015. Skeletal consequences of RANKL-blocking antibody (IK22-5) injections during growth: Mouse strain disparities and synergic effect with zoledronic acid. Bone 73:51–59. doi:10.1016/j.bone.2014.12.011

Li J-Y, Chassaing B, Tyagi AM, Vaccaro C, Luo T, Adams J, Darby TM, Weitzmann MN, Mulle JG, Gewirtz AT, Jones RM, Pacifici R. 2016. Sex steroid deficiency-associated bone loss is microbiota dependent and prevented by probiotics. J Clin Invest 126:2049–2063. doi:10.1172/JCI86062

Li J-Y, Tawfeek H, Bedi B, Yang X, Adams J, Gao KY, Zayzafoon M, Weitzmann MN, Pacifici R. 2011. Ovariectomy disregulates osteoblast and osteoclast formation through the T-cell receptor CD40 ligand. Proceedings of the National Academy of Sciences 108:768–773. doi:10.1073/pnas.1013492108

Li R, Boer CG, Oei L, Medina-Gomez C. 2021. The Gut Microbiome: a New Frontier in Musculoskeletal Research. Curr Osteoporos Rep 19:347–357. doi:10.1007/s11914-021-00675-x

Li T-H, Liu L, Hou Y-Y, Shen S-N, Wang T-T. 2019. C-type lectin receptor-mediated immune recognition and response of the microbiota in the gut. Gastroenterol Rep (Oxf) 7:312–321. doi:10.1093/gastro/goz028

Love MI, Huber W, Anders S. 2014. Moderated estimation of fold change and dispersion for RNA-seq data with DESeq2. Genome Biology 15. doi:10.1186/s13059-014-0550-8

Madel M-B, Ibáñez L, Ciucci T, Halper J, Rouleau M, Boutin A, Hue C, Duroux-Richard I, Apparailly F, Garchon H-J, Wakkach A, Blin-Wakkach C. 2020. Dissecting the phenotypic and functional heterogeneity of mouse inflammatory osteoclasts by the expression of Cx3cr1. eLife 9:e54493. doi:10.7554/eLife.54493

Madel M-B, Ibáñez L, Rouleau M, Wakkach A, Blin-Wakkach C. 2018. A Novel Reliable and Efficient Procedure for Purification of Mature Osteoclasts Allowing Functional Assays in Mouse Cells. Frontiers in Immunology 9. doi:10.3389/fimmu.2018.02567

Madel M-B, Ibáñez L, Wakkach A, de Vries TJ, Teti A, Apparailly F, Blin-Wakkach C. 2019. Immune Function and Diversity of Osteoclasts in Normal and Pathological Conditions. Frontiers in Immunology 10. doi:10.3389/fimmu.2019.01408

Merck E, Gaillard C, Gorman DM, Montero-Julian F, Durand I, Zurawski SM, Menetrier-Caux C, Carra G, Lebecque S, Trinchieri G, Bates EEM. 2004. OSCAR is an FcRgamma-associated receptor that is expressed by myeloid cells and is involved in antigen presentation and activation of human dendritic cells. Blood 104:1386–1395. doi:10.1182/blood-2004-03-0850

Mócsai A, Humphrey MB, Van Ziffle JAG, Hu Y, Burghardt A, Spusta SC, Majumdar S, Lanier LL, Lowell CA, Nakamura MC. 2004. The immunomodulatory adapter proteins DAP12 and Fc receptor gamma-chain (FcRgamma) regulate development of functional osteoclasts through the Syk tyrosine kinase. Proc Natl Acad Sci USA 101:6158–6163. doi:10.1073/pnas.0401602101

Moré MI, Swidsinski A. 2015. Saccharomyces boulardii CNCM I-745 supports regeneration of the intestinal microbiota after diarrheic dysbiosis - a review. Clin Exp Gastroenterol 8:237–255. doi:10.2147/CEG.S85574

Ochi S, Shinohara M, Sato K, Gober H-J, Koga T, Kodama T, Takai T, Miyasaka N, Takayanagi H. 2007. Pathological role of osteoclast costimulation in arthritis-induced bone loss. Proc Natl Acad Sci U S A 104:11394–11399. doi:10.1073/pnas.0701971104

Ohlsson C, Engdahl C, Fåk F, Andersson A, Windahl SH, Farman HH, Movérare-Skrtic S, Islander U, Sjögren K. 2014. Probiotics protect mice from ovariectomy-induced cortical bone loss. PLoS ONE 9:e92368. doi:10.1371/journal.pone.0092368

Pais P, Almeida V, Yilmaz M, Teixeira MC. 2020. Saccharomyces boulardii: What Makes It Tick as Successful Probiotic? J Fungi (Basel) 6. doi:10.3390/jof6020078

Park-Min K-H, Ji J-D, Antoniv T, Reid AC, Silver RB, Humphrey MB, Nakamura M, Ivashkiv LB. 2009. IL-10 suppresses calcium-mediated costimulation of receptor activator NF-kappa B signaling during human osteoclast differentiation by inhibiting TREM-2 expression. J Immunol 183:2444–2455. doi:10.4049/jimmunol.0804165

Patro R, Duggal G, Love MI, Irizarry RA, Kingsford C. 2017. Salmon provides fast and bias-aware quantification of transcript expression. Nature Methods 14:417–419. doi:10.1038/nmeth.4197

Pimentel H, Bray NL, Puente S, Melsted P, Pachter L. 2017. Differential analysis of RNA-seq incorporating quantification uncertainty. Nature Methods 14:687–690. doi:10.1038/nmeth.4324

Reyes C, Hitz M, Prieto-Alhambra D, Abrahamsen B. 2016. Risks and Benefits of Bisphosphonate Therapies. Journal of Cellular Biochemistry 117:20–28. doi:10.1002/jcb.25266

Rice PJ, Adams EL, Ozment-Skelton T, Gonzalez AJ, Goldman MP, Lockhart BE, Barker LA, Breuel KF, DePonti WK, Kalbfleisch JH, Ensley HE, Brown GD, Gordon S, Williams DL. 2005. Oral Delivery and Gastrointestinal Absorption of Soluble Glucans Stimulate Increased Resistance to Infectious Challenge. J Pharmacol Exp Ther 314:1079–1086. doi:10.1124/jpet.105.085415

Sancho D, Reis e Sousa C. 2012. Signaling by Myeloid C-Type Lectin Receptors in Immunity and Homeostasis. Annual Review of Immunology 30:491–529. doi:10.1146/annurev-immunol-031210-101352

Sato M, Sano H, Iwaki D, Kudo K, Konishi M, Takahashi H, Takahashi T, Imaizumi H, Asai Y, Kuroki Y. 2003. Direct binding of Toll-like receptor 2 to zymosan, and zymosan-induced NF-kappa B activation and TNF-alpha secretion are down-regulated by lung collectin surfactant protein A. J Immunol 171:417–425. doi:10.4049/jimmunol.171.1.417

Savva A, Roger T. 2013. Targeting Toll-Like Receptors: Promising Therapeutic Strategies for the Management of Sepsis-Associated Pathology and Infectious Diseases. Front Immunol 4. doi:10.3389/fimmu.2013.00387

Seeling M, Hillenhoff U, David JP, Schett G, Tuckermann J, Lux A, Nimmerjahn F. 2013. Inflammatory monocytes and Fcγ receptor IV on osteoclasts are critical for bone destruction during inflammatory arthritis in mice. PNAS 110:10729–10734. doi:10.1073/pnas.1301001110

Souza PPC, Lerner UH. 2019. Finding a Toll on the Route: The Fate of Osteoclast Progenitors After Toll-Like Receptor Activation. Frontiers in Immunology 10. doi:10.3389/fimmu.2019.01663

Takami M, Kim N, Rho J, Choi Y. 2002. Stimulation by toll-like receptors inhibits osteoclast differentiation. J Immunol 169:1516–1523. doi:10.4049/jimmunol.169.3.1516

Terciolo C, Dapoigny M, Andre F. 2019. Beneficial effects of Saccharomyces boulardii CNCM I-745 on clinical disorders associated with intestinal barrier disruption. Clin Exp Gastroenterol 12:67–82. doi:10.2147/CEG.S181590

Xu Z, Xie Z, Sun J, Huang S, Chen Y, Li C, Sun X, Xia B, Tian L, Guo C, Li F, Pi G. 2020. Gut Microbiome Reveals Specific Dysbiosis in Primary Osteoporosis. Front Cell Infect Microbiol 10:160. doi:10.3389/fcimb.2020.00160

Yahara Y, Barrientos T, Tang YJ, Puviindran V, Nadesan P, Zhang H, Gibson JR, Gregory SG, Diao Y, Xiang Y, Qadri YJ, Souma T, Shinohara ML, Alman BA. 2020. Erythromyeloid progenitors give rise to a population of osteoclasts that contribute to bone homeostasis and repair. Nat Cell Biol 22:49–59. doi:10.1038/s41556-019-0437-8

Yamasaki T, Ariyoshi W, Okinaga T, Adachi Y, Hosokawa R, Mochizuki S, Sakurai K, Nishihara T. 2014. The Dectin 1 Agonist Curdlan Regulates Osteoclastogenesis by Inhibiting Nuclear Factor of Activated T cells Cytoplasmic 1 (NFATc1) through Syk Kinase. J Biol Chem 289:19191–19203. doi:10.1074/jbc.M114.551416

Yu L, Zhao X-K, Cheng M-L, Yang G-Z, Wang B, Liu H-J, Hu Y-X, Zhu L-L, Zhang S, Xiao Z-W, Liu Y-M, Zhang B-F, Mu M. 2017. Saccharomyces boulardii Administration Changes Gut Microbiota and Attenuates D-Galactosamine-Induced Liver Injury. Sci Rep 7:1359. doi:10.1038/s41598-017-01271-9

Zaiss MM, Jones RM, Schett G, Pacifici R. 2019. The gut-bone axis: how bacterial metabolites bridge the distance. Journal of Clinical Investigation 129:3018–3028. doi:10.1172/JCI128521

Zhu X, Zhao Y, Jiang Y, Qin T, Chen J, Chu X, Yi Q, Gao S, Wang S. 2017. Dectin-1 signaling inhibits osteoclastogenesis via IL-33-induced inhibition of NFATc1. Oncotarget 8:53366–53374. doi:10.18632/oncotarget.18411

Zou W, Kitaura H, Reeve J, Long F, Tybulewicz VLJ, Shattil SJ, Ginsberg MH, Ross FP, Teitelbaum SL. 2007. Syk, c-Src, the αvβ3 integrin, and ITAM immunoreceptors, in concert, regulate osteoclastic bone resorption. Journal of Cell Biology 176:877–888. doi:10.1083/jcb.200611083

